# Forecasting Perception Before It Happens: Context-Specific Connectivity Patterns Predict Perceptual Outcomes

**DOI:** 10.1101/2025.03.06.641785

**Authors:** Parham Mostame, Guido Hesselmann, Richard Bido-Medina, Andreas Kleinschmidt, Sepideh Sadaghiani

## Abstract

The behavioral relevance of the static functional connectome is well-established, but the *real-time* relationship between ongoing connectome dynamics and behavior remains unclear. Because behavior shifts from moment to moment, an important question arises: Are changes in behavioral outcomes linked to fluctuations in task-specific or task-general processes like arousal? Task-specific versus task-general processes should manifest as distinct versus shared connectome patterns that predict moment-to-moment behavioral outcomes across tasks.

We analyzed fMRI data from three groups of healthy participants (total N=35), each performing one of three ambiguous perception tasks maximizing behavioral fluctuations: (1) Reporting faces or vase percepts on Rubin’s figure, (2) detecting near-random motion, or (3) identifying a near-threshold tone. Leveraging long inter-stimulus intervals (>20s), we examined how pre-stimulus connectome states influenced perception on a trial-by-trial basis.

Using SVM models, pre-stimulus connectome reliably predicted *upcoming* post-stimulus perception. Distinct, non-overlapping sets of task-specific connections supported prediction in each task, spanning task-relevant sensory networks as well as higher-order cognitive control networks. Only when predictive connections were aggregated over the well-known intrinsic connectivity networks (ICNs), did we observe partial overlap across tasks.

Our findings underscore the functional impact of ongoing connectome dynamics in shaping moment-to-moment behavior, and show that this impact is predominantly context-dependent.

## 2. Introductions

Brain activity is continuously ongoing, independent of external stimuli (Berger, 1930; Raichle, 2009, 2015). In the context of task-based studies, ongoing brain activity is often methodologically treated as noise, e.g. left unparameterized in regression analyses or averaged out in event-related analyses. However, in resting state studies ongoing brain activity has been recognized as functionally relevant (Raichle, 2015; Ramduny and Kelly, 2025). Importantly, a limited number of task-based studies suggest that ongoing brain activity and connectivity affect perception of consecutive stimuli on a *moment-to-moment* basis (cf. studies discussed below).

The moment-to-moment behavioral relevance of ongoing activity was first reported in electrophysiological recordings. Specifically, it has been shown that pre-stimulus activity levels correlate with the post-stimulus evoked activity in anesthetized cats (Arieli et al., 1996), and that pre-stimulus activity and connectivity predict visual threshold perception in monkeys (Supèr et al., 2003). Scalp EEG/MEG studies in awake humans have shown that perceptual outcomes across trials are linked to power of pre-stimulus oscillations (Linkenkaer-Hansen et al., 2004; Dijk et al., 2008; Keil et al., 2014; Kayser et al., 2016; Podvalny et al., 2019).

In fMRI research, the inherently slow dynamics of the hemodynamic response complicate efforts to distinguish behavioral effects driven by ongoing neural activity from those triggered by external stimuli. To address this, some perceptual fMRI studies have attempted to isolate ongoing activity by regressing out the average stimulus-evoked hemodynamic response. The residual fluctuations, particularly in the pre-stimulus period, are then interpreted as reflecting intrinsic neural processes. Such fluctuations have been linked to somatosensory perception (Boly et al., 2007), visual signal detection (Schölvinck et al., 2012), and object categorization (Wu et al., 2024).

Another line of fMRI research has leveraged distinctively long inter-stimulus intervals to more directly assess the moment-to-moment link between ongoing activity and perceptual outcomes. These studies have shown that the pre-stimulus fMRI signal amplitude can predict trial-by-trial motor speed (Thompson et al., 2013) and stimulus detection success (Sadaghiani et al., 2009; Goodale et al., 2021) in vigilance tasks. Such signals also predict moment-to-moment perceptual decisions including on a face/vase ambiguous stimulus (Rubin figure) (Hesselmann et al., 2008a) and a near-threshold random motion perception task (Hesselmann et al., 2008b). The long inter-trial periods in these studies allow for the examination of ongoing brain activity during tasks without the influence of evoked brain responses and their trial-to-trial variability, which cannot be regressed out.

Beyond ongoing *activity* discussed above, it has been established that dynamic large-scale *connectivity* during resting state is likewise of functional significance (Preti et al., 2017). These studies demonstrate associations between functional connectivity changes—often during task-free recordings—and behavioral performance measured separately (Vidaurre et al., 2017; Casorso et al., 2019; Eichenbaum et al., 2021; Jun et al., 2022). Such observations render it critical to investigate the moment-to-moment link of ongoing connectivity to perceptual outcomes *during* task performance. However, task-based studies are faced with the challenge of isolating the behavioral impact of ongoing from that of stimulus-driven connectivity fluctuations. Regression approaches have been proposed as one avenue to attempt this isolation (Ploner et al., 2010).

Only a few task-based connectivity studies have taken approaches that more directly separate stimulus-related from ongoing connectivity changes based on long inter-trial intervals. In one such study, we previously reported that detection of a near-threshold tone is preceded by greater connectivity in Cingulo-opercular and Default mode networks and lower connectivity in the Dorsal attention network (2015a). Further, we found that perceptual decisions are associated with pre-stimulus global network topology, specifically the degree to which canonical neurocognitive networks are segregated or conversely integrated. However, such fMRI studies employing long inter-trial intervals— crucial for isolating pre-stimulus connectivity from stimulus-evoked responses—are rare. Therefore, a key question has remained unanswered: Are the spontaneous connectivity fluctuations associated with perceptual outcomes largely the same or predominantly different across tasks?

Two major scenarios are possible. Firstly, the presumed moment-to-moment behavioral relevance of ongoing connectome dynamics could emerge from a universal set of connections across cognitive tasks, reflecting fluctuations in *task-common* processes. Spontaneous arousal fluctuations or attentional state shifts may be an example of such processes that exist independently of the context of the cognitive task. Chang and colleagues have shown that ongoing changes in alertness robustly relate to spontaneous changes in cognitive performance and spontaneous brain processes (Chang et al., 2016; Turchi et al., 2018). The instantaneous behavioral relevance of connectome dynamics may also manifest underlying fluctuations in the efficacy of cognitive control processes such as maintenance of task-set rules or attentional states (Rosenberg et al., 2015; Kucyi et al., 2017). For instance, waxing and waning of activity in the Cingulo-Opercular network could affect instantaneous vigilance (Goodale et al., 2021), therefore behavioral performance, as it plays a role in the maintenance of task-set rules (Sadaghiani and D’Esposito, 2015). Similarly, elevated activity in Default Mode intrinsic connectivity network (heretofore ICN) may be associated with lower cognitive performance across a wide range of datasets. If such task-general processes were the cause of ongoing behavioral fluctuations, they would emerge regardless of the cognitive context.

In the second scenario, the moment-to-moment changes of perceptual outcomes could be explained by context-dependent processes, manifested as changes in a set of unique connections in every task. Particularly, contextual differences could entail variability in sensory information (e.g. Visual or Auditory) or task rules and goals (e.g. decision-making as in two-alternative forced choice tasks or signal detection as in threshold detection tasks). Such context-dependent processes would likely engage connectivity changes within and among sensory systems as well as high order cognitive control networks. Such context-specific observations have been interpreted in the predictive coding theory of brain function where the ongoing processes act as priors for the upcoming perceptual processes (Friston, 2010; Hesselmann et al., 2010).

Finally, it can be conceived that there are substantial contributions to the association with perceptual outcomes from both task-common and task-specific connectivity fluctuations. Thus, investigating the task-specificity of the contribution of connections in the upcoming perceptual outcomes sheds light on what neurocognitive processes underlie the putative brain-to-behavior association of ongoing connectivity processes.

Answering the above-detailed question requires direct comparison of the role of spontaneous connectivity fluctuations across multiple distinct cognitive tasks. Here we first demonstrate that a machine learning model based on spontaneous pre-stimulus connectome dynamics (as measured in fMRI) can predict perception of subsequent ambiguous stimuli in three cognitively distinct tasks, in agreement with the predictive coding theory of the brain function (Friston, 2010; Hesselmann et al., 2010). Comparing the predictive connections across tasks, we then answer the critical question of whether a set of *task-common* or *task-specific* processes drives such a moment-to-moment association. Our findings at the level of individual connections and intrinsic connectivity networks (ICNs) speak for the latter; particularly involving connectivity within and among sensory modalities as well as cognitive control networks. These findings shed light on the connectivity processes that underlie the moment-to-moment changes of the behavior, in conjunction with advancing theoretical frameworks for studying connectomics.

## 3. Materials & Methods

Analysis code is provided at https://github.com/connectlab/predict_perception_Mostame2025. Time series of all atlas regions in continuous as well as epoched form are freely accessible at the Illinois Data Bank at: https://doi.org/10.13012/B2IDB-5233629_V1. Informed consent for sharing raw data was not obtained as part of the original studies for which the data was collected.

### 3.1. Data & participants

We used fMRI recordings of a total of 35 healthy participants (17 females; 19-30 age range) in three dissimilar tasks: Rubin, Motion detection, and Auditory vigilance (Fig. 1). Participant groups differed between tasks but were recruited by the same procedures from the same community and underwent an identical imaging sequence at the same MRI scanner. Participants had no history of neurological or psychiatric disorders, were right-handed, and gave written informed consent to participate. A.K. had ethics approval for these studies. The Rubin task comprised twelve participants (age range: 20–29 years; 9 females), the Motion detection task twelve participants (age range: 19–30 years, 6 females) and the Auditory vigilance task eleven participants (age range: 19–30 years; 2 females).

**Fig. 1.**
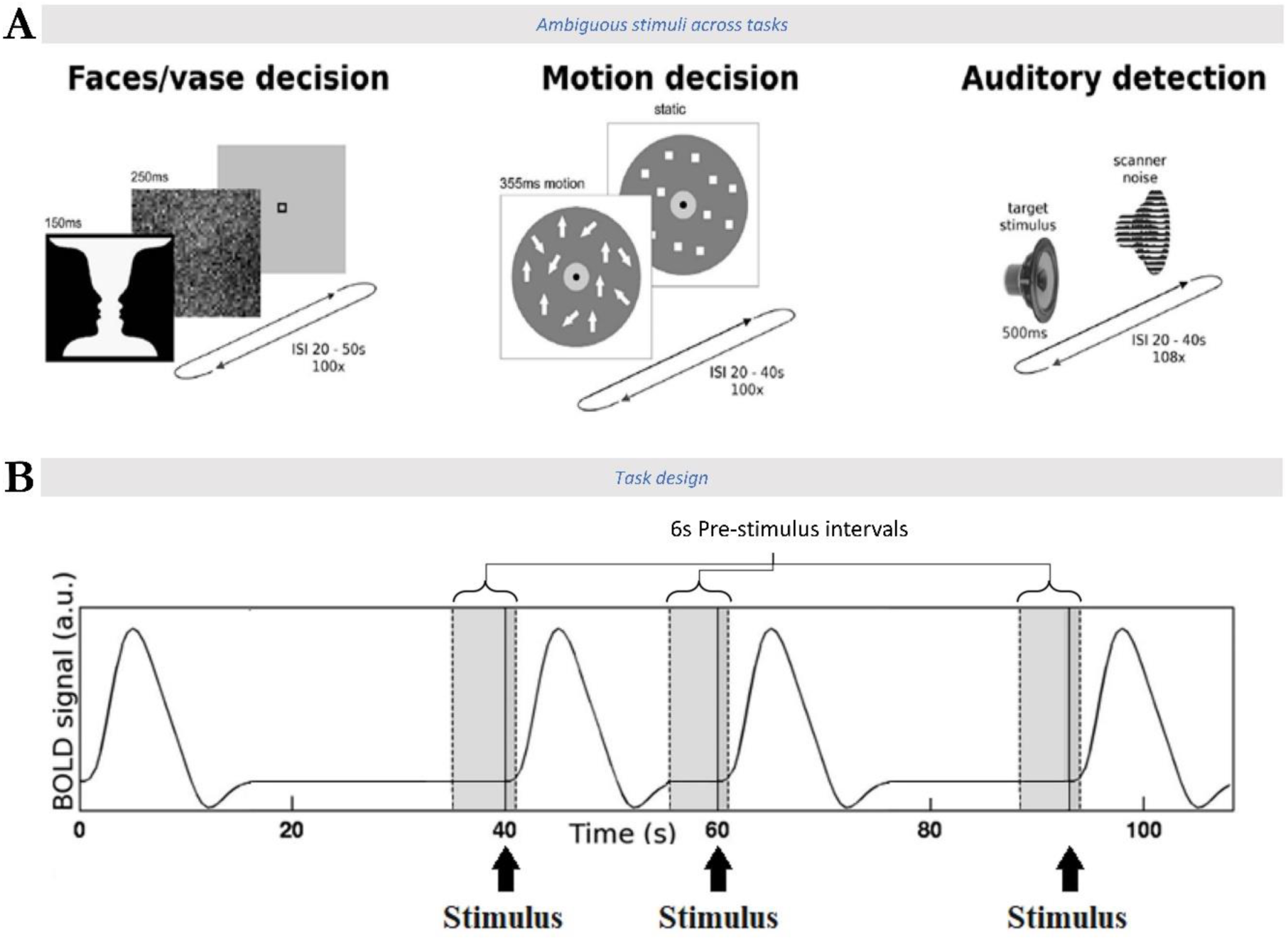
Task information across all datasets. A) Task paradigms for Rubin, Motion detection, and Auditory vigilance tasks from left to right. The near-threshold stimuli and very long inter-stimulus-intervals guaranteed inter-trial variability in the perceptual responses of the participants. B) 6 seconds of the pre-stimulus interval (−5 to +1s relative to stimulus onset) was used for prediction of perceptual outcomes and for network analysis of predictive connections. Figures were obtained from (Sadaghiani et al., 2010a) and modified with permission.

In each task, participants were presented with the same threshold-level or ambiguous stimulus across all trials (plus catch trials; cf. below), which however resulted in different percepts across trials. Specifically, the stimulus in each task was either ambiguous by nature (Rubin) or presented at the participant-specific perceptual threshold (Motion and Auditory). This experimental design allowed the investigation of perceptual fluctuations due to ongoing changes of connectome configurations. Participants’ perceptual response to the identical stimulus in each task determined whether a trial is labeled as (analytic) target or non-target for the purpose of machine learning model training. Importantly, these tasks were uniquely designed to have a very long inter-stimulus interval (>∼20s) allowing for the study of ongoing connectome processes free of evoked hemodynamic responses (Sadaghiani et al., 2010a). In the following, we briefly describe each task.

#### 3.1.1. Statistical power

The central question in this work is whether the pre-stimulus state of the connectome affects subsequent perception in a *context-dependent* manner. Therefore, the most critical statistical analysis is the comparison across cognitive contexts, specifically the one-way ANOVA depicted in Figure 2C. A one-way ANOVA (fixed effects) with a total of 35 subjects over 3 independent groups has 80% power (p<0.05 uncorrected) to detect an effect size of *f*=0.55. In other words, even prior to multiple comparisons correction (FDR), the connection-wise difference across tasks needs to be large to be detected (Cohen, 1969). Therefore, our current sample will not be able to detect small to medium-sized differences across tasks. However, any significant effects observed after correction would indicate the presence of large cross-task differences in the perceptual impact of pre-stimulus connectome states.

**Fig. 2.**
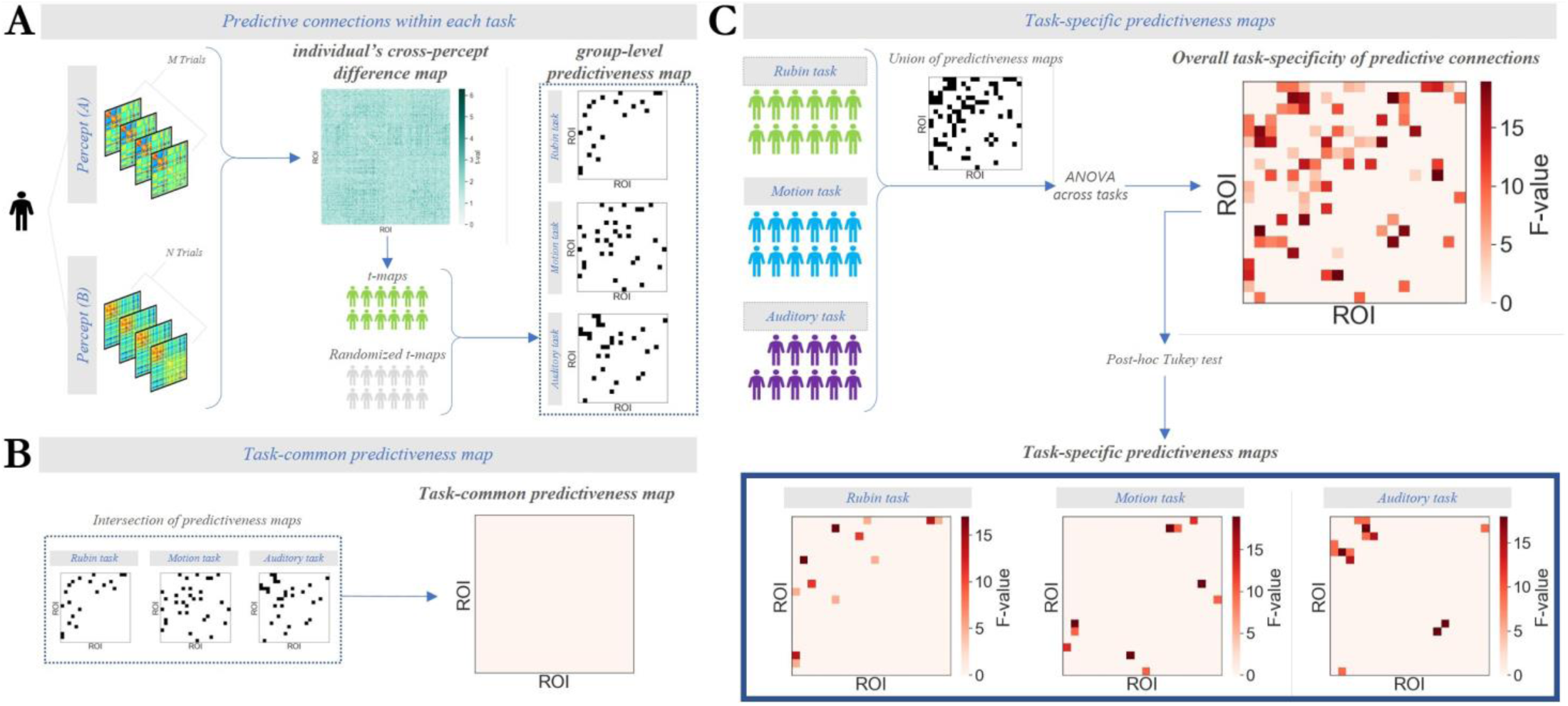
Statistical analysis pipeline for extracting task-common and task-specific predictive connections. A) Extraction of predictive connections within each task is depicted. The difference of pre-stimulus connectivity strength across the two percepts (left) was quantified at each connection per participant, resulting in an individual cross-percept difference map for each participant (center). In each task, a connection-wise group-level paired t-test was then performed between the maps from all participants and participant-specific surrogate maps (randomized t-maps; cf. methods). This step resulted in a group-level t-map indicating predictive connections in each task. These three maps were binarized into group-level predictiveness maps (right) B) A task-common predictiveness map was extracted from the intersection of group-level predictiveness maps across all tasks. No connection was predictive in more than one task, resulting in an empty task-common predictiveness map. C) Every connection that appeared in the group-level predictiveness map of at least one task (cf. 2A right), i.e. union across tasks, entered a connection-wise one-way ANOVA. The ANOVA comprised task as a factor with three levels and participants’ cross-percept difference in each task (cf. individuals’ cross-percept difference map, 2A center) as samples with random effects (Benjamini Hochberg FDR-corrected for number of connections). Then, post-hoc Tukey tests identified connections that were more predictive in a particular task than at least one other task, resulting in three task-specific predictiveness maps.

#### 3.1.2. Rubin task

This data is described in full detail elsewhere (Hesselmann et al., 2008a). In brief, across two 25-minutes sessions participants were presented with the ambiguous Face/Vase Rubin figure over a total of 90 trials (plus 10 catch trials with the upside-down image and not analyzed in this study) (Fig. 1). Participants were instructed to report via two buttons whether they perceived “*Faces*” or “*Vase*”, where the face percept was chosen as the analytic target in our machine learning model. The Rubin figure was presented for 150ms, immediately followed by a 250ms noise mask with equivalent contrast. The ISIs were extracted from a gamma-like distribution spanning 20s to 50s (i.e. with most ISIs being from the short end). The original study showed that the proportions of responses (58% *Faces* averaged across participants) and the reaction times (*Faces*: 812±72ms; *Vase*: 842±75ms, no significant difference) were similar across *Faces* and *Vase* responses.

#### 3.1.3. Motion task

The motion task dataset was originally introduced elsewhere (Hesselmann et al., 2008b). In brief, 500 white squares of size 0.2 inch were randomly distributed on a dark gray annulus of size 23 inches (Fig. 1). Participants were instructed to maintain gaze at the center of the annulus marked by a blue rectangle (1 inch) surrounded by a 3-inches light gray circle (Fig. 1B). Most squares moved in a random-walk fashion, while a small percentage moved in a coherent fashion up- or down-ward. The percentage for the latter was extracted individually for each participant prior to the main experiment as the participant’s threshold for perceiving coherent movement on half of the trials. Participants were asked to respond whether they perceived a coherent (“*Coherent*”) or random (*“Random”*) movement on each trial, where the *Coherent* percept was chosen as the analytic target in our machine learning model. Over two 25-minutes sessions, there were a total of 60 trials at threshold coherence level plus 40 catch trials at very low or very high coherence (not analyzed in this study). Stimuli had the length of 355ms and were separated by an ISI of 20s to 40s extracted from a uniform random distribution. The original study showed that although there was no considerable difference between the proportion of *Coherent* and *Random* percepts (57% *Coherent* on average), participants responded significantly faster (*t*_11_ = 4.03, p<0.01) for the coherent (1160ms) than the random perceptual response (1324ms).

#### 3.1.4. Auditory task

This data has been previously published elsewhere (Sadaghiani et al., 2009). Briefly, the auditory stimulus was a 500ms noise burst with its frequency band modulated at 2 Hz, presented at each participant’s auditory threshold (Fig. 1). Individuals’ auditory threshold was estimated by a staircase procedure at the beginning of the scan. Participants were blind-folded and asked to report by a key press as soon as they heard the auditory stimulus (responses within 1.5s of presentation were considered as hits). The percept was labeled as a “*Hit*” when the participants detected the tone, and a “*Miss*” when the tone was not perceived. Note that the hit condition was chosen as the analytic target in our machine learning model of this task. All participants performed two 20-minutes runs while some conducted an additional third run (time permitting). Each run consisted of 36 trials at threshold volume and four catch trials with a supra-threshold volume, and with a random ISIs between 20s and 40s. Before each run, the stimulus was played a few times at a supra-threshold level so that participants could re-memorize the stimulus over the scanner background noise. The near-threshold stimulus was perceived in slightly more than half of the trials (62%) with a reaction time of 788±102ms. False alarms were rare overall (4.5 per subject, with an interquartile range of 6.9). Particularly, only a negligible fraction of these false alarms (a median of 2 with an interquartile range of 3.2 per session and across subjects) fell within the 20 seconds preceding the upcoming stimulus onset.

Stimuli in all three tasks were presented at unpredictable intervals. However, the auditory task differs from the two visual tasks in an important way. In the auditory task, stimulus occurrence was certain while perceptual detection was uncertain. In contrast, perceptual detection in the visual tasks was certain, but stimulus identity was ambiguous. This conceptual difference increases the divergence of cognitive context across tasks and allows us to test whether ongoing connectome patterns predict perceptual outcomes in a task-common or task-specific manner. In other words, we intentionally avoided closely matched paradigms to enable a direct assessment of context dependence.

### 3.2. Image data acquisition

MRI data for all tasks were acquired at the same three-Tesla whole-body scanner (Tim-Trio; Siemens) with identical sequences. Anatomical images were recorded with a T1-weighted magnetization-prepared rapid acquisition gradient echo sequence [176 slices, repetition time (TR) = 2300 ms, echo time (TE) = 4.18 ms, filed of view (FOV) = 256 x 256, voxel size 1 x 1 x 1 mm). Functional imaging used a T2*-weighted gradient-echo, echo-planar-imaging sequence (25 slices, TR = 1500 ms, TE = 30 ms, FOV= 192 x 192, voxel size 3 x 3 x 3 mm, interslice gap 20%).

### 3.3. Data preprocessing

Standard preprocessing was applied as part of the original three studies (Hesselmann et al., 2008a, 2008b; Sadaghiani et al., 2015a) using statistical parametric mapping (SPM5, Wellcome Department of Imaging Neuroscience, UK; www.fil.ion.ucl.ac.uk), comprising rigid-body realignment, coregistration, and normalization to MNI stereotactic space. No spatial smoothing was applied.

Head motion as quantified by mean framewise displacement (FD, (Power et al., 2012)) applied to all pre-stimulus timepoints that entered the analyses. FD was smaller than 0.5 for all individuals in each task. Further, there was little to no difference in head motion between the two prestimulus conditions in all tasks. Particularly, mean FD difference between the percepts was equal to 0.001 for Rubin task (*t_11_*=1, *p*=0.34); 0.006 for Motion task (*t_11_*=1.04, *p*=0.32), and 0.008 for Auditory task (*t_11_*=2.27, *p*=.05).

The following indicates the preprocessing steps of all three tasks.

### 3.4. Estimation of pre-stimulus connectivity

To estimate whole-brain connectivity, we adopted the brain parcellation from (Power et al., 2011) with 264 regions. This parcellation has the advantage of being defined jointly under consideration of task-related co-activations and resting state co-fluctuations. Further, as we previously applied this same parcellation in our prior investigation of the Auditory detection dataset (Sadaghiani et al., 2015a), keeping the choice consistent allowed us to directly compare our outcomes with the previous study. The BOLD signal was averaged across all voxels within a 6mm-radius sphere centered at the coordinates provided for each of 264 regions by (Power et al., 2011) and further constrained by a gray matter mask. The BOLD signal of each ROI was high-pass filtered with a 128-second cutoff to remove low-frequency drifts and regressed against nuisance signals, including head motion parameters, motion outliers (DVARS), and compartment regressors (white matter, gray matter, and cerebrospinal fluid). FIR regressors were also included to account for task-related activity, modeled using a finite impulse response (FIR) basis set with 24 regressors per condition (1.5 s time bins), covering a 36s window around stimulus onset. For the auditory task, regressor conditions included false alarms in addition to Hit and Miss. Brain regions lacking coverage in any participant of a given task were excluded from machine learning in that task (Rubin: 0, Motion: 28, and Auditory: 26 ROIs), enabling downstream cross-task comparison of connectomes. For the cross-task comparisons of predictive connections, all these regions were excluded so that only regions available in all tasks were retained. For this analysis, we further excluded the 28 regions of the Power-Petersen atlas (Power et al., 2011) that were not assigned to any of the atlas’s 13 brain networks (see Table S4 in Supplementary materials).

Framewise connectome dynamics (Esfahlani et al., 2020) was extracted from the pre-stimulus interval of all trials of all participants. The pre-stimulus interval was defined as the interval of −5 to +1 seconds relative to stimulus onset. Note that the hemodynamic fMRI signal during the first second of post-stimulus interval does not include any stimulus-related information as shown in previous studies using current datasets (Sadaghiani et al., 2010). Further, this period is free of hemodynamic response to the previous trial due to the long inter-trial intervals.

### 3.5. Machine learning models for prediction of perceptual outcomes within each task

To depict the presence of a moment-to-moment association between ongoing connectome dynamics and behavior, we adopted a predictive analysis within each task. Particularly, we set out to establish predictability of perceptual outcomes in each task (*Faces* vs. *Vase*, *Coherent* vs. *Random*, and *Hit* vs. *Miss*, respectively) using a Support Vector Machine (SVM) model that is trained on the pre-stimulus connectivity data as defined above separately in each task.

Once pre-stimulus connectome patterns were calculated (cf. 3.4), the upper triangular section of each connectome (i.e. matrix) was vectorized and pooled across trials and participants. Cross-validation was performed using a 20-fold approach across all trials, where each fold allocated 95% of trials for training and 5% for testing. Particularly in each fold, training and test data consisted of approximately 738 and 39 trials in Rubin task, 655 and 34 trials in Motion task, and 687 and 36 trials in Auditory task (pooled across subjects and percepts).

As in our prior work (Sadaghiani et al., 2015a), the approach of pooling data over participants was necessary due to the low number of trials in each participant. The low trial number is a direct effect of the design choice of having very long inter-trial intervals for the study of evoked response-free inter-stimulus intervals. While pooling adds substantial participant-specific variance (noise) to the model, any success of this approach would suggest that pre-stimulus connectome states exert an impact on subsequent perception that is common across the participants of our experiment. Given that the data are pooled across subjects, we do not claim generalizability of the prediction across individuals.

To avoid data leakage, we ensured that connectome frames of the same trials would never split across train and test data. For each fold, the train and test data were normalized with respect to the mean and standard deviation of the features in the train data. Additionally, all preprocessing steps within the predictive pipeline including normalization, feature selection, and hyperparameter optimization, were performed using only the training data of each fold and subsequently evaluated on the held-out test data. Further, selected trials across data folds were stratified over percepts and participants, which ensured uniform distribution of participants and percepts across folds.

F-score feature selection was performed prior to model training separately for each task. To capture FC variability *within* pre-stimulus timepoints, feature selection was performed separately at every timepoint (−5 to +1s) pooled over participants. Then, for each feature (i.e., connection), the highest *F*-score across all pre-stimulus timepoints of trials was selected. Eventually, a grid search approach was adopted to maximize model performance across variable number of selected features (N = 25, 50, 100, 200, 400, 800 connections out of all connections).

Once the features were identified, an SVM model with a Radial Basis Function (RBF) kernel was trained on the preprocessed connectivity data of each task. We used a secondary grid search to find the best model parameters including gamma (8 points between −5 and l in log scale) and C (8 points between −1 and 4 in log scale). Model performance was evaluated by different metrics (averaged across data folds) including accuracy, balanced accuracy, and area under the curve of the Receiver Operating Characteristic curve (AUC-ROC) (Fig. 3). All reported performance metrics were computed exclusively on held-out test data and averaged across repetitions. The confusion matrix of predictive models of all tasks are included in the supplementary materials (Fig. S1). Finally, evaluation metrics were tested against a null distribution generated from randomly shuffled analytic target labels of the data (R=200).

**Fig. 3.**
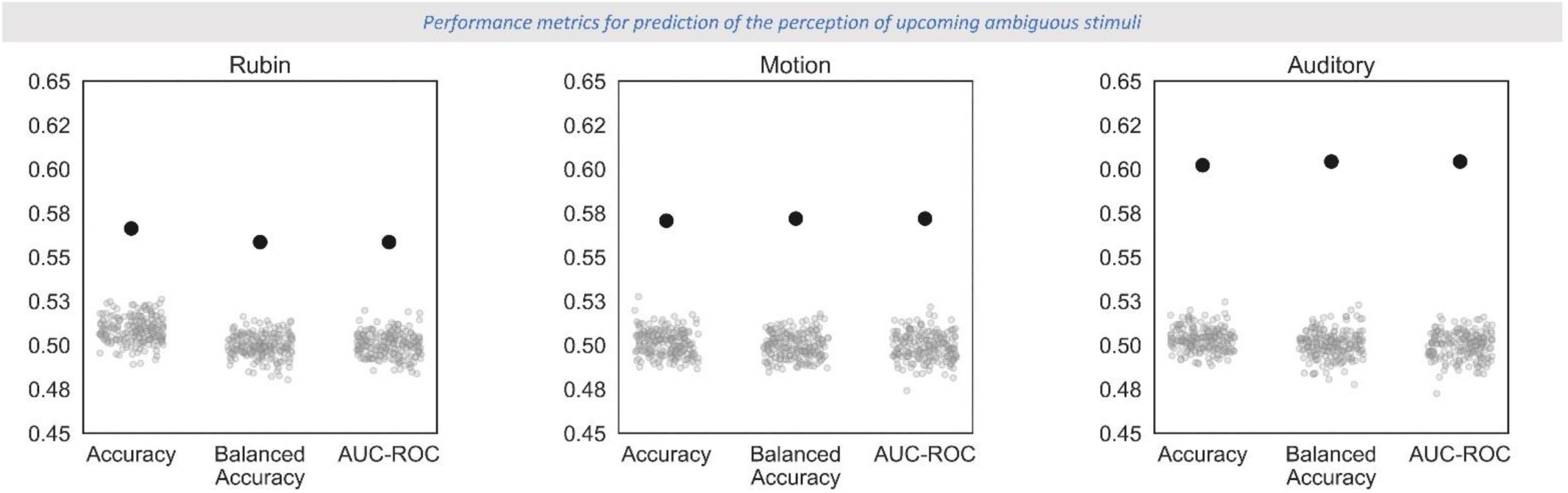
Model performance across tasks. For each task, the plot depicts the respective SVM model’s ability to predict trial-by-trial perceptual outcomes based on pre-stimulus connectivity. Each subplot illustrates model performance using Accuracy, Balanced Accuracy, and AUC-ROC metrics in comparison to 200 surrogate samples generated from randomly shuffled analytic target labels. Our models exceed chance level across all metrics for all tasks, showing that pre-stimulus connectome dynamics can predict post-stimulus perceptual outcomes. This finding demonstrates the moment-to-moment behavioral relevance of spontaneous connectome reconfigurations.

### 3.6. Predictive connections across tasks

After establishing that pre-stimulus connectome states can predict perception in all three tasks, we assessed whether this predictability arises from the same or different connections across tasks. The aforementioned SVM feature selection for *within*-task predictions was performed at the *group-level*. The downstream comparison of predictive connections *across* the three tasks with independent participant cohorts, however, requires an individual-level feature selection to account for cross-cohort biases. Thus, our entire *cross-task* comparison analysis is based on an alternative *individual-level* feature selection approach described in the following.

#### Individual cross-percept difference maps

Within each participant and separately for each task, pre-stimulus timepoints (−5 to +1s) from all trials were separated into two sets according to the two perceptual outcomes. For every connection in the entire connectome, connectivity values were compared between the two sets using a two-sample *t*-test at each connection. This analysis calculates a single difference matrix per task and participant (see Fig. 2A, center). Note that these maps do not serve statistical evaluation in themselves, but rather provide a measure of subject-wise cross-percept difference (power of mean difference across percepts when normalized by their corresponding variance) for further analysis.

#### Group-level predictiveness maps

All *individual cross-percept difference maps* in each task were then tested against participant-specific null maps using a group-level paired *t*-test in a connection-wise manner. Each participant’s null map corresponded to the connection-wise mean of 500 surrogate maps. Each surrogate map of a participant was generated by randomly shuffling the participant’s original individual cross-percept difference map over connections. This design ensures that percept-related effects are quantified relative to that individual’s own connectivity fingerprint rather than against zero across subjects, minimizing cohort-specific influences. Additionally, including age and sex as covariates in the cross-task ANOVA analysis yielded virtually identical results (Fig. S5). Together, these steps ensure that the cross-task comparisons are unlikely to be driven by cross-cohort differences.

The group analysis using paired *t*-test across subjects resulted in a *t*-map. The *t*-map was then thresholded at uncorrected p<0.01 to obtain a group-level binary mask (Fig. 2A, right). Note that we did not apply multiple comparisons correction over connections because the group-level *t*-maps are not generated for statistical evaluation in themselves. Instead, they serve further analysis of *task-common* and *task-specific predictiveness maps* as detailed below.

#### Task-common predictiveness map

In this step, we sought to answer which connections, if any, are predictive of perceptual outcome in a *context-<u>in</u>dependent* manner. To this end, we identified connections that were consistently predictive across all tasks by extracting the intersection of the connections that were present in the *group-level predictive maps* across all three tasks (Fig. 2B). Because no connections fell within this intersection (cf. Results), no multiple comparisons correction (i.e. further restriction of connections) was necessary.

#### Task-specific predictiveness maps

In this step, we sought to answer which connections, if any, are predictive of perceptual outcome in a *context-dependent* manner. First, we created the union of *group-level predictiveness maps* across the three tasks for further analysis. In other words, any connection that did not pass the group-level *t-*test in at least one task was discarded from this analysis. Then, task-specificity of each of the remaining connections was tested independently using an ANOVA test. For each connection, predictiveness values extracted from participant-specific *t*-maps of all tasks were compared in a one-way ANOVA with three levels corresponding to the three tasks. This resulted in an *F* value per connection indicating the extent of task-specificity in the “predictiveness” of that connection (top panel in Fig. 2C). The significance of *F*-values was corrected for multiple comparison using Benjamini Hochberg FDR correction at *q*=0.05.

Following the ANOVA, post-hoc Tukey tests were performed to identify connections that were more predictive in a particular task compared to both other tasks (Family-Wise Error corrected for 3 comparisons at *p*=0.05). Overall, this analysis yielded three *task-specific predictiveness maps* (bottom panel in Fig. 2C, resulting in Fig. 4). Each of these maps comprises the original *F*-values but for a *subset* of the connections from the ANOVA. Additionally in each map, the direction of change in connectivity values across percepts was extracted based on raw differences of connectivity values across the percepts, where the positive direction corresponds to the analytic target label (as color-coded in Fig. 4). Note that we replicated the ANOVA analysis with the inclusion of demographic covariates to ensure that our cross-task comparisons are not driven by cross-cohort age and sex differences (see Fig. S5).

**Fig. 4.**
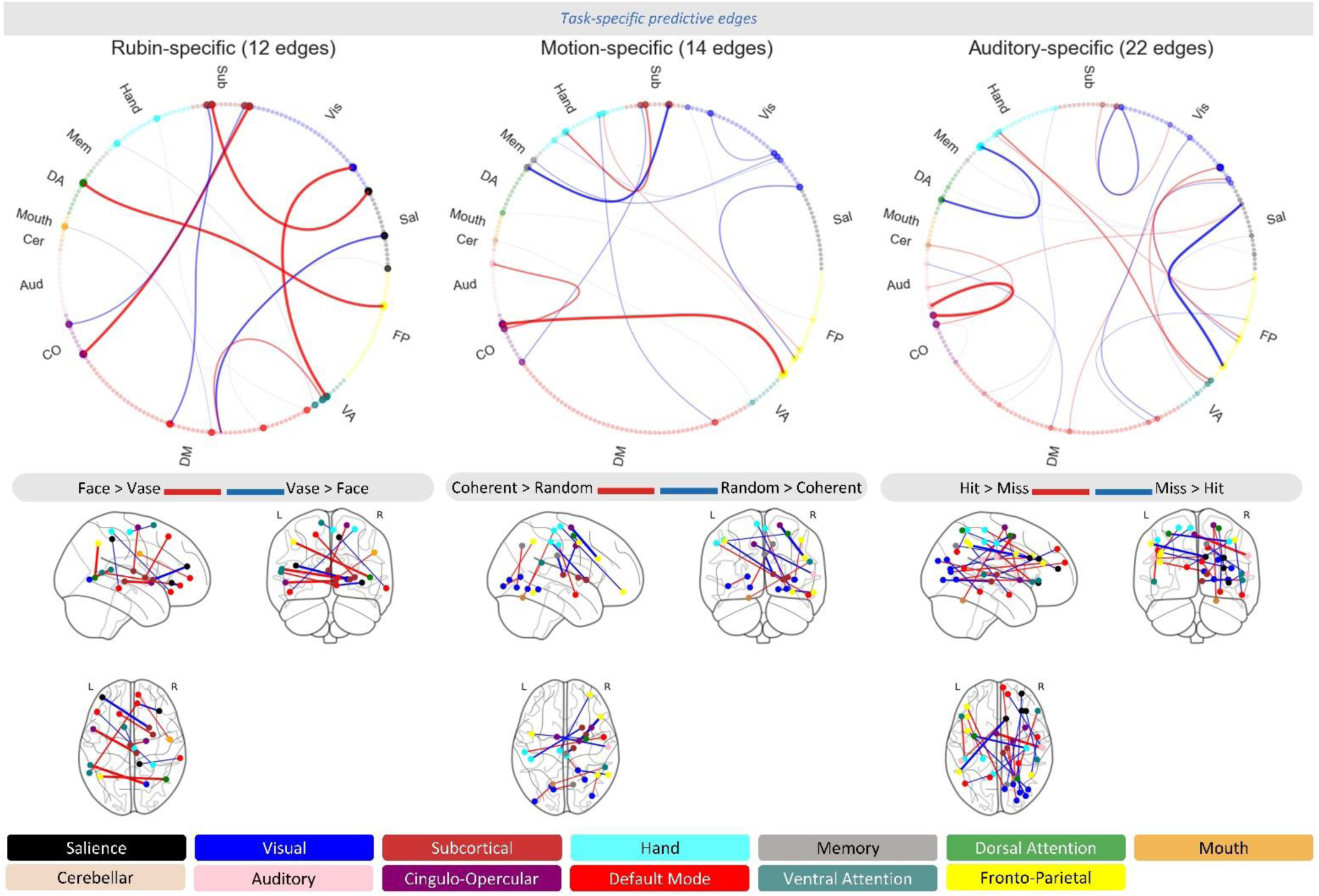
Graph view of the predictive connections in Rubin faces-vase decision, Motion detection, and Auditory vigilance tasks from left to right. Each of the visualized connections was more predictive of the perceptual outcome in the respective task than both other tasks. Connections associated with perception in any task (cf. Fig 2A) were compared across tasks by a one-way ANOVA followed by post-hoc Tukey tests. Multiple comparison correction was applied to the ANOVA’s F values (Benjamini-Hochberg FDR correction at q<0.05) as well as the Tukey tests (FWE correction at p<0.05). Red or blue indicates which of the two percepts was associated with pre-stimulus increase in the connectivity strength of the respective connection. The line thickness corresponds to F-values, which reflect the extent of task-specificity of the connection. For network label abbreviations, see methods. The MNI coordinates of each of the regions involved (in counterclockwise order) are provided in Supplementary materials Tables S1 to S3 for the three tasks.

### 3.7. ICN-level predictiveness across tasks

To interpret results in relation to well-known neurocognitive ICNs, we used ICN labels provided by Power et al. (2011) for all regions of the parcellation. These ICNs include (somatosensory) *Hand*, (somatosensory) *Mouth*, Cingulo-Opercular (*CO*), Auditory (*Aud*), Default Mode (*DM*), Memory (*Mem*), Ventral Attention (*VA*), Visual (*Vis*), Fronto-Parietal (*FP*), Salience (*Sal*), Sub-Cortical (*Sub*, consisting of thalamus and basal ganglia), Cerebellar (*Cer*), and Dorsal Attention (*DA*) networks. An additional set of regions that could not be assigned to any ICN by Power et al. was labeled as N/A and excluded from our predictive edge detection analyses.

This analysis was descriptive and aimed at aiding interpretation of the connection-wise analyses detailed above. Specifically, we quantified ICN-level connectome differences associated with perceptual outcomes of each task as follows. We summed connection-wise *F*-values of the *task-specific predictiveness maps* over all ICN pairs, separately for connections in which stronger pre-stimulus connectivity predicted the analytic target or non-target percept, respectively. Then, the sum was divided by the total number of connections across the respective ICN pair (resulting in Fig. 5). ICN-level task-commonality was then qualitatively characterized by the intersection of ICN-level predictiveness maps across all tasks (illustrated in Fig. 6).

**Fig. 5.**
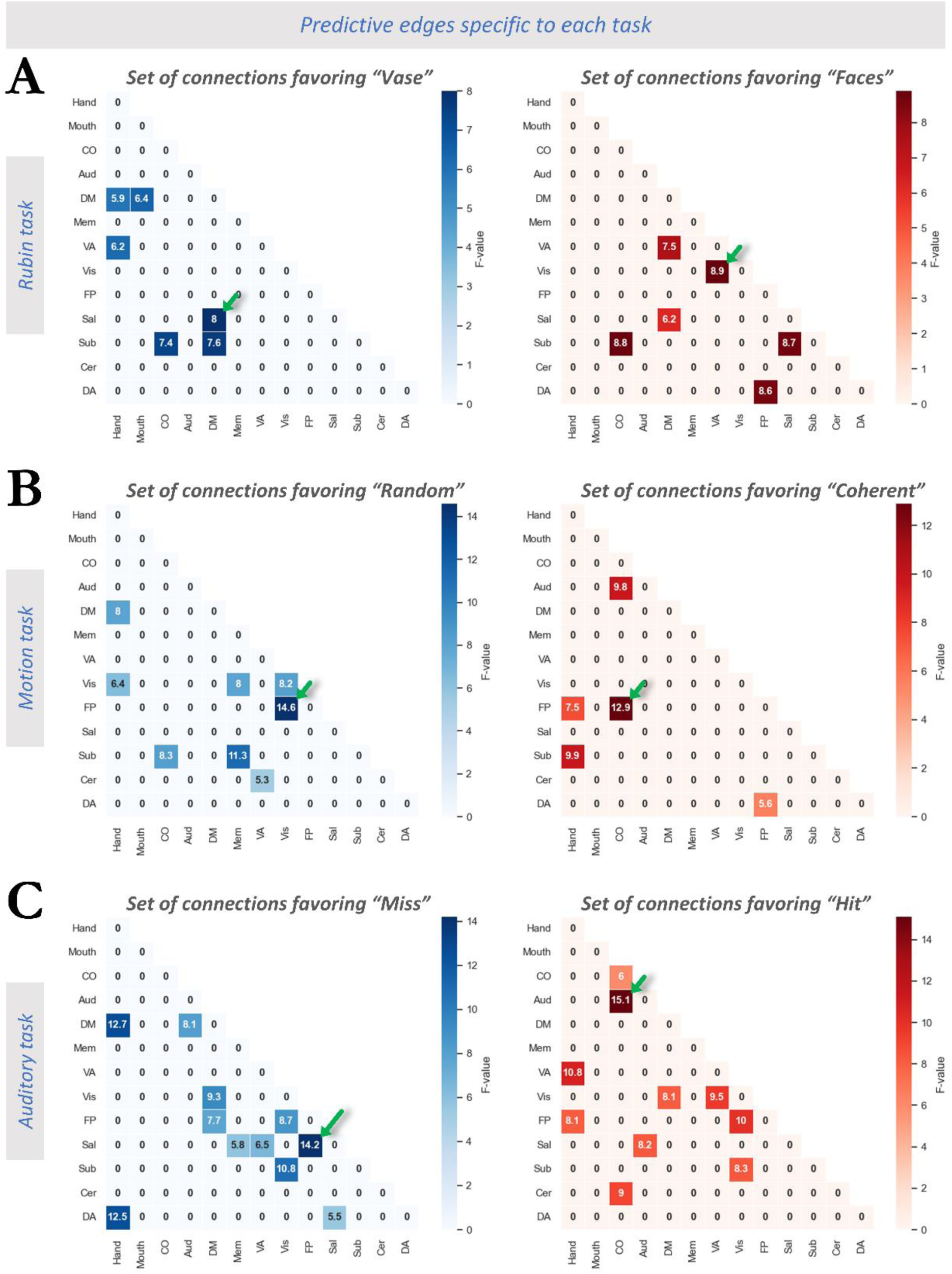
Task-specific predictive connections fall within specific ICNs. Each matrix illustrates the co-engagement of each pair of canonical ICNs in the respective task-specific predictiveness map (cf. Fig 4). In other words, the F-values (i.e. strength of task-specificity) of the connections in Fig. 4 are aggregated within pairs of ICNs. The left (blue heatmap) and right (red heatmap) panels represent ICN pairs in which connectivity was higher prior to the analytic non-target and analytic target percepts, respectively. The strongest effect in each panel as marked by a green arrow is briefly described here, with additional effects discussed in the main text. A) Rubin task. Face percepts are more likely when the task-relevant Visual (Vis) network is more strongly connected to the Ventral Attention (VA) network (right side), and the Default Mode (DM) network is more dis-engaged from the Salience (Sal) network (left). B) Motion detection task. Coherent motion percepts are associated with disengagement of the Fronto-Parietal (FP) network from the Visual network (left) and its simultaneous engagement with other cognitive control networks including the Cingulo-Opercular (CO) network (right). C) Auditory vigilance task. Hits are associated with co-engagement of the task-relevant Auditory (Aud) network with the Cingulo-Opercular network (right) and the disengagement of Salience and Fronto-Parietal networks (left).

**Fig. 6.**
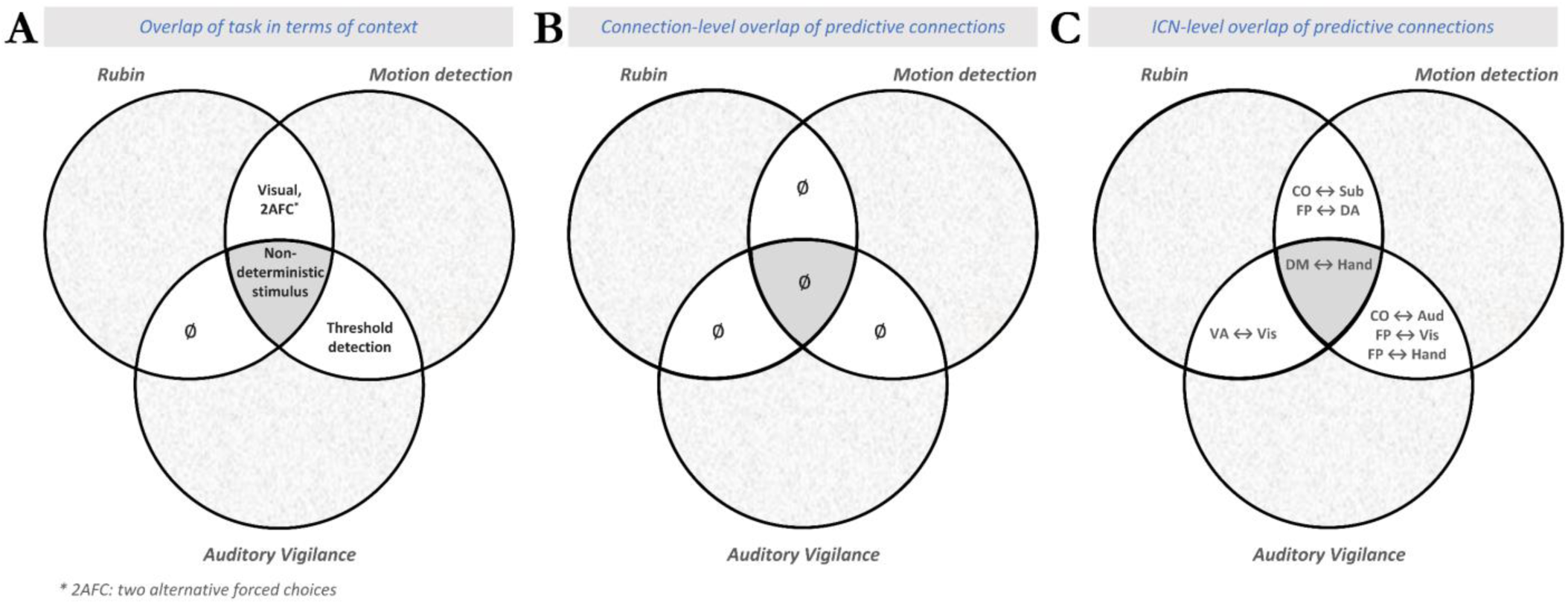
Overlap of the three tasks in context, predictive connections, and predictive ICNs. Each task presents a unique context emerging from the interplay of several task characteristics. While some of these characteristics are unique to each task (hashed gray areas), this figure focuses on the overlapping aspects. A) Ven diagram illustrating the overlap of the three tasks in terms of characteristics such as sensory modality. The dark gray area in the center represents the ambiguous nature shared across all tasks. The overlap of every pair of tasks is depicted as white areas. B) similar to (A) but for overlap of predictive features across tasks, particularly at the level of connections. C) similar to (B) but at a coarser granularity of ICN network interactions. These observations suggest that despite the task-specific predictive connections, upcoming perception associates with multiple task-common network-level interactions including both sensory systems and high order cognitive control networks. 2AFC: two-alternative forced choice paradigm.

### 3.7 Global network topology

We investigated changes in global network topology, specifically the Modularity Index (Newman, 2006), in relation to consecutive perceptual outcomes (Fig. S7). This approach identifies neural correlates of moment-to-moment perceptual changes in terms of pre-stimulus fluctuations in the global architecture of the connectome rather than in individual connections (Zalesky et al., 2010). The cross-percept comparison of modularity index was conducted as follows.

We first computed the average connectomes of all trials for each of two percepts of every individual. Next, we calculated the modularity index (Q) of each participant and percept as follows. We used the Power-Petersen atlas to define community structure (Power et al., 2011), where each canonical ICN was considered a community. For each connectome, we quantified modularity by assessing within-community connection strength relative to the expected strength under a null model. Specifically, for each community, we computed the total within-community edge weight (WD) and normalized it by the total edge weight of the network (m), applying a resolution parameter (γ=1) to account for module size effects. Eventually, a paired *t*-test was conducted separately for each task to compare modularity values between the two perceptual conditions (Fig. S7).

## 4. Results

### 4.1. Model performance in prediction of perceptual outcomes in each task

The SVM models predicted the perceptual outcome better than chance in all tasks. Fig. 3 shows the evaluation metrics averaged across all cross-validation folds, including the null metrics. For Rubin faces-vase decision, Motion detection, and Auditory vigilance tasks, model accuracy was 0.57, 0.57, and 0.60, respectively, significantly exceeding chance level (z score = 8.1, 9.5, 15.4). Similarly, the balanced accuracy and AUC metrics were equal to 0.56, 0.57, 0.60 (z score = 8.6, 10.8, 13.8), and 0.56, 0.57, 0.60 (z score = 8.6, 10.8, 13.8) for the three tasks, respectively. The corresponding confusion matrices are shown in Fig. S1. Moreover, the group-average pre-stimulus connectome organization of each percept and the corresponding cross-percept connectivity differences are included in the supplementary materials for each task (Fig. S2-S4).

### 4.2. Spatially characterizing predictive connections

Next, we aimed to answer the central question of whether the connectivity fluctuations associated with moment-to-moment perception are common across tasks or are task-specific instead. To answer this question, we compared the predictive features across tasks both at the level of individual connections and canonical ICNs.

#### 4.2.1. Connection-level cross-task comparisons

For each task, a binary *group-level predictiveness map* was constructed as a connection-wise paired *t*-test. Specifically, the group-level test compared each participant’s observed *cross-percept difference map* to the participant-specific surrogate (spatially randomized) maps (Fig 2A). Considering the intersection of these *group-level predictiveness maps* over all pairs of tasks (Fig. 2B), we did not find any overlapping connections even in a maximally lenient approach, i.e. even when group-level predictiveness maps were *not* corrected for multiple comparisons. In other words, we did not find any connections in which pre-stimulus connectivity fluctuations were associated with perceptual outcomes in more than one task (conceptually illustrated in Fig. 6B).

Given the lack of overlap, we next assessed whether each of the predictive connections is associated with perceptual outcome specifically in one task more so than the other tasks (Fig. 2C). In the union of the three *group-level predictiveness maps*, a one-way ANOVA of factor ‘task’ was applied to the participant-specific *cross-percept difference maps* (for replication under inclusion of demographic factors see Fig. S5). Connections identified as task-specific in this analysis entered pairwise post-hoc tests to identify connections more predictive in one than both other tasks. In the ensuing *task-specific predictiveness maps,* we found 10, 15, and 23 predictive connections for Rubin, Motion, and Auditory tasks, respectively (Fig. 4). The Power-Petersen ROIs involving these connections are listed for each task in Tables S1 to S3.

All three *task-specific predictiveness maps* spanned both sensory networks as well as higher-order control networks. Pre-stimulus connectivity between the task-relevant sensory (Visual or Auditory) networks and various higher-order networks was associated with perception in all tasks. For example, in the case of the Auditory vigilance task higher connectivity between auditory cortex and the CO network (specifically dorsal anterior cingulate cortex) was observed before successful sound detection. In the case of the Rubin task, high pre-stimulus connectivity between a Visual network region in the right ventral stream and the Ventral Attention network was predictive of perceiving *Faces* rather than a *Vase*. Interestingly, in the visual Motion detection task high connectivity to Visual regions resulted in perceiving the dot motion as *Random*, i.e. missing the subtle motion coherence. Contrarily, a connection of the Dorsal Attention network, specifically the right Frontal Eye Field (FEF), to a frontal region of the Fronto-Parietal network (i.e. posterior middle frontal gyrus) was stronger prior to perceiving motion *Coherence*. This observation suggests that the pre-stimulus connectivity state of the Dorsal Attention network (critical in motion perception (Shadlen and Newsome, 1996)), biases perception toward coherence, while heightened connectivity of the Visual network may promote sensitivity to lower-level visual features, leading to a perception of randomness.

#### 4.2.2. The distribution of task-specific predictive connections across canonical ICNs

Given the observed task-specificity of predictive connections, we next asked whether these connections occurred among the same or different ICNs across tasks. The following approach aims at aiding network-level interpretation of the results from the ANOVA and post-hoc tests from Fig. 4. To this end, we aggregated the ANOVA-derived measure of task-specificity within each pair of canonical ICNs. This aggregation was performed across the predictive connections identified for each task (Fig. 5). The resulting ICN-level task-specificity was then compared across tasks (Fig. 6C). Essentially, this descriptive analysis quantifies to what degree the connections shown on the three subplots of Fig. 4 occur within the same ICN pairs. Note that while we focus the following discussion on ICN-pairs with particularly strong outcomes, these effects are distributed widely, spanning multiple ICNs.

Connections in which high pre-stimulus connectivity biased towards perceiving a *Vase* are shown in Fig. 5A left. Here, the DM network stands out with particularly high connectivity to Sal, Sub (i.e., thalamus and basal ganglia), motor-Hand and motor-Mouth networks. Additionally, high connection density occurred between CO and Sub networks. In contrast, connections in which high pre-stimulus connectivity preceded *Faces* percepts (Fig. 5A right) occurred most strongly between VA and two other networks, specifically Vis and DM. Other ICN-pairs included Sub to both CO and Sal.

Fig. 5B left represents connections in which high pre-stimulus connectivity preceded the *Random* percept. Here, the Vis network stood out, with high within-network (Vis-Vis) connectivity, as well as connectivity to motor-Hand, Mem, and FP networks. Other ICN pairs included Sub to CO and Mem. Conversely, connections in which high pre-stimulus connectivity preceded *Coherent* motion perception (Fig. 5B right) occurred between the DA network (involved in motion perception) and the FP network. The FP network also stood out, with high connectivity to CO and motor-Hand networks.

Auditory detection performance depended on connectivity of the task-relevant Aud network, as well as CO, Sal, and DM cognitive control networks (Fig. 5C). In particular, co-engagement of the Aud network with both CO and Sal networks facilitated perception of the tone (5C right). Conversely, co-engagement of the DM network with several networks, specifically Aud, Vis, motor-Hand, and FP, biased towards *Misses* (5C left). Beyond the latter network interaction (FP-DM), co-engagement of FP network with Vis and Sal likewise preceded *Misses*.

#### 4.2.3. Summary of Results

Figure 6 summarizes our observations. Despite the partial *contextual* overlap of some task pairs (Fig. 6A), the individual predictive connections did not converge across any combination of the tasks (Fig. 6B, which summarizes lack of overlap in Fig. 4). However, at the level of network interactions (Fig. 6C, which summarizes Fig. 5), multiple network pairs emerged that were associated with the upcoming perception in more than one task.

Converging across all three tasks, we found the co-engagement of Motor-Hand and DM networks biasing towards perception of the analytic non-targets (*Vase*, *Random*, and *Miss* percepts in Rubin, Motion detection, and Auditory vigilance tasks, respectively). Conversely, co-activation of Motor-Hand with a different higher-order network, specifically the FP network, preceded the opposite perceptual outcome in both detection tasks, i.e. Auditory and Motion, toward *Hit* and *Coherent* percepts, respectively.

We also found other network interactions converging between two (but not three) tasks. For instance, in both visual tasks, upcoming perception was linked to co-engagement of CO and Sub as well as DA and FP networks. However, as expected from the lack of overlapping connections, the regions in these ICN pairs differed across tasks. For DA-FP connectivity, for instance, the connection between *posterior* regions of the two networks biased towards *Faces* over *Vase* percepts in the Rubin task. Conversely, the connectivity of *anterior* (frontal) regions of these networks supported *Coherent* motion detection.

### 4.3. Regional activity versus distributed connectivity

We sought to ensure that the significance of pre-stimulus brain states for subsequent perception indeed includes impact from connectivity *across* region-pairs as opposed to solely activity *within* regions. This is critical because our FC measure reflects short-lived connectivity processes that estimate instantaneous co-activation levels of region pairs. To this end, we replicated our predictive models jointly using regional activations *and* cross-regional FC values as model features. Firstly, we did not observe an increase in model performance in any task (see Supplementary Materials). Secondly, although both regional activations and cross-region FCs were chosen during feature selection in all tasks, the percentage of features selected among all available activity features was significantly lower than the corresponding percentage of FC features (p<0.011 across all tasks). This outcome supports the functional relevance of cross-region connectivity in predicting upcoming perception over and above regional activity alone (Fig. S6).

## 5. Global network topology

We assessed whether large-scale changes in the spatial organization of the connectome, can affect upcoming perception beyond the observed cross-areal FC changes. To this end, we quantified and compared global modularity for the connectome states preceding the two percepts in each task (Fig. S7). For the Auditory vigilance task, modularity was higher prior to *Hits* compared to *Misses* (paired t-test; *t_10_* = 2.5; p = 0.032). This observation replicates our previous work in the Auditory task (Sadaghiani et al., 2015a), despite using a different analysis pipeline and FC measure. Interestingly however, we did not find a significant difference in global modularity across percepts in Rubin and Motion detection tasks.

## 6. Discussion

We established a moment-to-moment link between ongoing connectome dynamics and perception. Specifically, we showed that ongoing connectome dynamics can predict the perception of upcoming ambiguous stimuli using machine learning models. One framework to understand this observation is the predictive coding theory of brain function, suggesting that ongoing connectome dynamics act as prior knowledge influencing our perception of the environment (Friston, 2010; Hesselmann et al., 2010). Notably, perceptual outcomes were associated with connectome changes involving both sensory networks as well as higher-order cognitive networks, though in a task-specific manner. This observed task-specificity of the brain-to-behavior association aligns with the idea that the brain hosts a repertoire of cognitive architectures, each represented by a distinct connectome pattern (Petersen and Sporns, 2015). From this perspective, each cognitive “architecture”—that is, intrinsic connectivity pattern—modulates perception in a way that reflects its functional role with the specific cognitive context.

We sought to disambiguate two scenarios in which connectome dynamics may influence perception – either through context-dependent neural representations or via general cognitive control mechanisms. Context-dependent modulation is expected to diverge across tasks differing in sensory modality, task rules etc., whereas general cognitive control mechanisms such as arousal should influence perception regardless of the specific task paradigm. In agreement with context-dependent modulation, several earlier studies have reported that pre-stimulus ongoing “activity” in task-relevant regions profoundly impacts the perception of upcoming ambiguous stimuli (Hesselmann et al., 2008b, 2008a; Sadaghiani et al., 2009; Schölvinck et al., 2012). However, the impact of pre-stimulus activity on perception has also been observed in higher-order cognitive control networks (Boly et al., 2007; Sadaghiani et al., 2010b; Wu et al., 2024), though assessing the degree to which such an impact may be shared across cognitive contexts would require a direct comparison across tasks.

Turning to pre-stimulus connectivity, our study offers a direct cross-task comparison that sheds light on the specificity of connectome-perception relationships. The finding that predictive connections vary by task (see Fig. 4) supports the scenario of context-dependent modulation. However, when examined at the network level, we observed that the fluctuations in connectivity across certain cognitive control networks consistently influenced perceptual outcomes across multiple tasks (see Fig. 5-6), suggesting an additional contribution of general cognitive control mechanisms.

We propose that two interconnected processes may underlie the observed task-specific predictive connections. First, ongoing fluctuations in connectivity involving task-relevant sensory systems may serve as a form of prior representation, guiding perception of upcoming stimuli. For example, connectivity involving a ventral region of the Vis network in the Rubin task, the Frontal Eye Fields (FEF) regions of DA network in the Motion detection task, and the Superior Temporal Gyrus (STG) and insular regions of Aud network in the Auditory vigilance task were each associated with their corresponding consecutive perceptual outcomes. Second, the dynamic state of cognitive control networks and their interactions with sensory and other control networks also influenced perception. For instance, connectivity between the FP network and both the Vis network and other cognitive control networks (DA and CO) was linked to subsequent perceptual outcomes.

Although we did not find task-common connections, investigating the task-commonality at the level of ICNs revealed that the Motor-Hand and DM networks consistently served as predictive features. This effect may reflect biases in neural representations, particularly fluctuations in the motor system’s readiness potential (Deecke, 1987), given that all tasks required manual responses via button presses. Alternatively, the effect may relate to the suppression of somatosensory modalities due to the antagonistic integration of multimodal sensory information during visual or auditory processing (Clements et al., 2022). Interestingly, however, connectivity between the Hand to FP networks predicted successful detection in both Auditory and Motion detection tasks—an effect opposite to what would be expected under the sensory suppression hypothesis. This observation suggests that executive control exerted by the FP network (Dosenbach et al., 2007; Seeley et al., 2007; Menon and D’Esposito, 2022) over motor cortex may facilitate the detection of near-threshold stimuli when decisions are manually reported.

One consideration in interpreting the absence of task-common connections is whether nuisance regression may have attenuated widespread cognitive state effects, such as arousal or attention. While the goal of compartment regression is typically the removal of non-neuronal shared variance (e.g., physiological and motion-related components), it is indeed important to consider that the global gray signal contains widespread neural activity as well (Schölvinck et al., 2010). While we cannot exclude the possibility that such widespread neural activity contains signals tracking attention and arousal, it should be noted that such signals have spatially characteristic maps (Rosenberg et al., *Nat Neurosci* 2016, Chang et al., PNAS 2016). They are most pronounced in specific brain networks that, while widespread, make up a relatively limited proportions of the brain’s gray matter. As such, global averaging with areas that do not carry attention or arousal signals is expected to dampen the representation of the latter in the global gray signal. However, since the balance of artifacts vs. neuromodulatory effects in the global signal might vary across participants and scans depending on (e.g.) the amount of head motion in a given scan, further studies may be needed to clarify the influence of global signal regression. Regarding task-common processes, we note that while we did not observe any individual edges that were predictive across all three tasks, there was partial convergence at the ICN level for specific task pairs (Fig. 6), consistent with the idea that large-scale processes related to attention/control may contribute but do so in a context-dependent way rather than via a universal edge-level signature.

Beyond connectivity changes between specific regions or networks, the overall topological configuration of the connectome as a whole-brain graph also holds functional significance (Preti et al., 2017). Yet, few studies have explored how spontaneous fluctuations in connectome topology relate to behavioral variability. Global network-based topology metrics, such as modularity, which reflects the balance between segregation and integration, have been proposed as key contributors to cognitive function (Zalesky et al., 2010; Shine and Poldrack, 2018). Sadaghiani et al. (2015b) previously reported that a highly modular connectome prior to detecting near-threshold auditory tones was associated with stimulus detection. We replicated this finding in the Auditory task (Fig. S7), underscoring the importance of prestimulus network topology beyond local connectivity. Interestingly however, no impact from modularity was observed for the other two tasks. This observation suggests that this global topological feature may exert a stronger influence when the task demands sustained vigilance. Vigilance is regulated by diffuse neuromodulatory neurotransmitter systems that broadly innervate cortical networks. Consequently, spontaneous fluctuations in vigilance are expected to manifest as widespread changes in neural activity and functional connectivity. Supporting this account, we previously reported peristimulus activity differences across percepts that were broadly distributed across numerous higher-order ICNs in the vigilance-dependent auditory task (Sadaghiani et al., 2009), while they were spatially restricted to task-relevant sensory areas in the Motion and Rubin tasks (Hesselmann et al., 2008b, 2008a). Consistent with these findings, the present study shows that prestimulus connectivity differences involved a substantially larger number of edges in the auditory task compared with the other tasks (Fig. 4). Given this spatially diffuse pattern, it is plausible that vigilance-related fluctuations are reflected at the level of global network topology. In this context, higher modularity may reflect greater segregation of large-scale networks, supporting sustained maintenance of task-relevant states and facilitating detection of near-threshold stimuli, whereas tasks driven by more transient perceptual decisions may rely less on such global organization.

### 6.1. Limitations and Future Directions

One limitation of the present study is that each task was conducted in a separate group of participants, which constrains the extent to which cross-task differences can be attributed exclusively to task context. While a within-participant, multi-task design would have been ideal, the present work leverages very rare existing datasets that uniquely share long inter-stimulus intervals, allowing isolation of pre-stimulus connectivity from stimulus-evoked responses. To mitigate subject-specific influences, our analysis focused on within-subject cross-percept differences evaluated relative to subject-specific null distributions. However, while this approach accounts for subject-specific connectome fingerprints, it cannot fully eliminate the possibility that cohort-level factors contributed to the observed effects. In particular, once individual connectivity fingerprints are minimized, remaining cross-task differences may reflect characteristics shared within a group but differing across groups. Although incorporating demographic variables where available revealed no significant influence (Fig. S5), other unmeasured factors—such as differences in domain-general strategies, attentional style, or processing dynamics—may still contribute to the apparent task specificity of predictive connectivity patterns. For example, systematic differences in how participants approach perceptual uncertainty or maintain task goals across cohorts could influence pre-stimulus network configurations in a way that mimics context-dependent effects. Therefore, the present findings should be interpreted as providing initial evidence for a predominantly context-dependent relationship between ongoing connectivity and perception, while future studies employing within-subject, multi-task designs will be critical to rule out potential latent group-level influences.

The sample size was further limited to sensitivity for large effect sizes only. Therefore, additional small to medium effects may have been missed. Additionally, the relatively low number of trials per participant, necessitated by the long inter-stimulus intervals required to capture unobscured pre-stimulus connectome states, imposed constraints on the performance of the machine learning models. Nevertheless, it is noteworthy that our models performed significantly better than chance across all tasks.

## 7. Conclusions

We showed that pre-stimulus large-scale connectome patterns can predict perceptual outcomes across three distinct tasks, suggesting that the influence of ongoing connectome dynamics on subsequent cognitive processes is a widespread phenomenon. Notably, no specific connection was predictive across multiple tasks; rather, the predictive connections were task-specific. This task-specificity indicates that a particular cognitive processes was associated with a unique connectome instantiation among the diverse repertoire of possible connectome states (Petersen and Sporns, 2015). At a broader level, some overlap emerged at the scale of ICNs, where perceptually predictive connectivity showed partial similarity across task pairs. Overall, our findings underscore the functional relevance of ongoing connectome states for moment-to-moment behavior, predominantly in a context-dependent manner.

## 8. Fundings & acknowledgements

SS was supported by NIH/NIMH grant R01MH116226.

## References

Arieli A, Sterkin A, Grinvald A, Aertsen A (1996) Dynamics of ongoing activity: explanation of the large variability in evoked cortical responses. Science 273:1868–1871.

Berger H (1930) Ueber das Elektrenkephalogramm des Menschen. [Electrocephalography in man.]. J Für Psychol Neurol 40:160–179.

Boly M, Balteau E, Schnakers C, Degueldre C, Moonen G, Luxen A, Phillips C, Peigneux P, Maquet P, Laureys S (2007) Baseline brain activity fluctuations predict somatosensory perception in humans. Proc Natl Acad Sci U S A 104:12187–12192.

Casorso J, Kong X, Chi W, Van De Ville D, Yeo BTT, Liégeois R (2019) Dynamic mode decomposition of resting-state and task fMRI. NeuroImage 194:42–54.

Chang C, Leopold DA, Schölvinck ML, Mandelkow H, Picchioni D, Liu X, Ye FQ, Turchi JN, Duyn JH (2016) Tracking brain arousal fluctuations with fMRI. Proc Natl Acad Sci 113:4518–4523.

Clements GM, Gyurkovics M, Low KA, Beck DM, Fabiani M, Gratton G (2022) Dynamics of alpha suppression and enhancement may be related to resource competition in cross-modal cortical regions. NeuroImage 252:119048.

Cohen J (1969) Statistical power analysis for the behavioral sciences. New York: Academic Press.

Deecke L (1987) Bereitschaftspotential as an indicator of movement preparation in supplementary motor area and motor cortex. Ciba Found Symp 132:231–250.

Dijk H van, Schoffelen J-M, Oostenveld R, Jensen O (2008) Prestimulus Oscillatory Activity in the Alpha Band Predicts Visual Discrimination Ability. J Neurosci 28:1816–1823.

Dosenbach NUF, Fair DA, Miezin FM, Cohen AL, Wenger KK, Dosenbach RAT, Fox MD, Snyder AZ, Vincent JL, Raichle ME, Schlaggar BL, Petersen SE (2007) Distinct brain networks for adaptive and stable task control in humans. Proc Natl Acad Sci 104:11073–11078.

Eichenbaum A, Pappas I, Lurie D, Cohen JR, D’Esposito M (2021) Differential contributions of static and time-varying functional connectivity to human behavior. Netw Neurosci 5:145–165.

Esfahlani FZ, Jo Y, Faskowitz J, Byrge L, Kennedy DP, Sporns O, Betzel RF (2020) High-amplitude cofluctuations in cortical activity drive functional connectivity. Proc Natl Acad Sci 117:28393–28401.

Friston K (2010) The free-energy principle: a unified brain theory? Nat Rev Neurosci 11:127–138.

Goodale SE, Ahmed N, Zhao C, de Zwart JA, Özbay PS, Picchioni D, Duyn J, Englot DJ, Morgan VL, Chang C (2021) fMRI-based detection of alertness predicts behavioral response variability Koyejo S, Büchel C, eds. eLife 10:e62376.

Hesselmann G, Kell CA, Eger E, Kleinschmidt A (2008a) Spontaneous local variations in ongoing neural activity bias perceptual decisions. Proc Natl Acad Sci 105:10984–10989.

Hesselmann G, Kell CA, Kleinschmidt A (2008b) Ongoing Activity Fluctuations in hMT+ Bias the Perception of Coherent Visual Motion. J Neurosci 28:14481–14485.

Hesselmann G, Sadaghiani S, Friston KJ, Kleinschmidt A (2010) Predictive Coding or Evidence Accumulation? False Inference and Neuronal Fluctuations. PLOS ONE 5:e9926.

Jun S, Alderson TH, Altmann A, Sadaghiani S (2022) Dynamic trajectories of connectome state transitions are heritable. NeuroImage 256:119274.

Kayser SJ, McNair SW, Kayser C (2016) Prestimulus influences on auditory perception from sensory representations and decision processes. Proc Natl Acad Sci 113:4842–4847.

Keil J, Müller N, Hartmann T, Weisz N (2014) Prestimulus Beta Power and Phase Synchrony Influence the Sound-Induced Flash Illusion. Cereb Cortex 24:1278–1288.

Kucyi A, Hove MJ, Esterman M, Hutchison RM, Valera EM (2017) Dynamic Brain Network Correlates of Spontaneous Fluctuations in Attention. Cereb Cortex N Y N 1991 27:1831–1840.

Linkenkaer-Hansen K, Nikulin VV, Palva S, Ilmoniemi RJ, Palva JM (2004) Prestimulus Oscillations Enhance Psychophysical Performance in Humans. J Neurosci 24:10186–10190.

Menon V, D’Esposito M (2022) The role of PFC networks in cognitive control and executive function. Neuropsychopharmacology 47:90–103.

Newman MEJ (2006) Modularity and community structure in networks. Proc Natl Acad Sci 103:8577–8582.

Petersen SE, Sporns O (2015) Brain Networks and Cognitive Architectures. Neuron 88:207–219.

Ploner M, Lee MC, Wiech K, Bingel U, Tracey I (2010) Prestimulus functional connectivity determines pain perception in humans. Proc Natl Acad Sci 107:355–360.

Podvalny E, Flounders MW, King LE, Holroyd T, He BJ (2019) A dual role of prestimulus spontaneous neural activity in visual object recognition. Nat Commun 10:3910.

Power JD, Barnes KA, Snyder AZ, Schlaggar BL, Petersen SE (2012) Spurious but systematic correlations in functional connectivity MRI networks arise from subject motion. Neuroimage 59:2142–2154.

Power JD, Cohen AL, Nelson SM, Wig GS, Barnes KA, Church JA, Vogel AC, Laumann TO, Miezin FM, Schlaggar BL, Petersen SE (2011) Functional network organization of the human brain. Neuron 72:665–678.

Preti MG, Bolton TA, Van De Ville D (2017) The dynamic functional connectome: State-of-the-art and perspectives. NeuroImage 160:41–54.

Raichle ME (2009) A Paradigm Shift in Functional Brain Imaging. J Neurosci 29:12729–12734.

Raichle ME (2015) The restless brain: how intrinsic activity organizes brain function. Philos Trans R Soc B Biol Sci 370:20140172.

Ramduny J, Kelly C (2025) Connectome-based fingerprinting: reproducibility, precision, and behavioral prediction. Neuropsychopharmacology 50:114–123.

Rosenberg MD, Finn ES, Constable RT, Chun MM (2015) Predicting moment-to-moment attentional state. NeuroImage 114:249–256.

Sadaghiani S, D’Esposito M (2015) Functional Characterization of the Cingulo-Opercular Network in the Maintenance of Tonic Alertness. Cereb Cortex 25:2763–2773.

Sadaghiani S, Hesselmann G, Friston K, Kleinschmidt A (2010a) The relation of ongoing brain activity, evoked neural responses, and cognition. Front Syst Neurosci 4 Available at: https://www.frontiersin.org/articles/10.3389/fnsys.2010.00020 [Accessed August 23, 2022].

Sadaghiani S, Hesselmann G, Friston K, Kleinschmidt A (2010b) The relation of ongoing brain activity, evoked neural responses, and cognition. Front Syst Neurosci 4 Available at: https://www.frontiersin.org/articles/10.3389/fnsys.2010.00020 [Accessed August 23, 2022].

Sadaghiani S, Hesselmann G, Kleinschmidt A (2009) Distributed and Antagonistic Contributions of Ongoing Activity Fluctuations to Auditory Stimulus Detection. J Neurosci 29:13410–13417.

Sadaghiani S, Poline J-B, Kleinschmidt A, D’Esposito M (2015a) Ongoing dynamics in large-scale functional connectivity predict perception. Proc Natl Acad Sci 112:8463–8468.

Sadaghiani S, Poline J-B, Kleinschmidt A, D’Esposito M (2015b) Ongoing dynamics in large-scale functional connectivity predict perception. Proc Natl Acad Sci 112:8463–8468.

Schölvinck, M. L., Maier, A., Ye, F. Q., Duyn, J. H., & Leopold, D. A. (2010). Neural basis of global resting-state fMRI activity. Proceedings of the National Academy of Sciences, 107(22), 10238–10243

Schölvinck ML, Friston KJ, Rees G (2012) The influence of spontaneous activity on stimulus processing in primary visual cortex. NeuroImage 59:2700–2708.

Seeley WW, Menon V, Schatzberg AF, Keller J, Glover GH, Kenna H, Reiss AL, Greicius MD (2007) Dissociable Intrinsic Connectivity Networks for Salience Processing and Executive Control. J Neurosci 27:2349–2356.

Shadlen MN, Newsome WT (1996) Motion perception: seeing and deciding. Proc Natl Acad Sci 93:628–633.

Shine JM, Poldrack RA (2018) Principles of dynamic network reconfiguration across diverse brain states. NeuroImage 180:396–405.

Supèr H, van der Togt C, Spekreijse H, Lamme VAF (2003) Internal State of Monkey Primary Visual Cortex (V1) Predicts Figure–Ground Perception. J Neurosci 23:3407–3414.

Thompson GJ, Magnuson ME, Merritt MD, Schwarb H, Pan W-J, McKinley A, Tripp LD, Schumacher EH, Keilholz SD (2013) Short-time windows of correlation between large-scale functional brain networks predict vigilance intraindividually and interindividually. Hum Brain Mapp 34:3280–3298.

Turchi J, Chang C, Ye FQ, Russ BE, Yu DK, Cortes CR, Monosov IE, Duyn JH, Leopold DA (2018) The Basal Forebrain Regulates Global Resting-State fMRI Fluctuations. Neuron 97:940–952.e4.

Vidaurre D, Smith SM, Woolrich MW (2017) Brain network dynamics are hierarchically organized in time. Proc Natl Acad Sci 114:12827–12832.

Wu Y, Podvalny E, Levinson M, He BJ (2024) Network mechanisms of ongoing brain activity’s influence on conscious visual perception. Nat Commun 15:5720.

Zalesky A, Fornito A, Bullmore ET (2010) Network-based statistic: Identifying differences in brain networks. NeuroImage 53:1197–1207.

