## Supplementary Materials for "Forecasting Perception Before It Happens: Context-Specific Connectivity Patterns Predict Perceptual Outcomes"

### Predicting perceptual outcomes from pre-stimulus connectivity

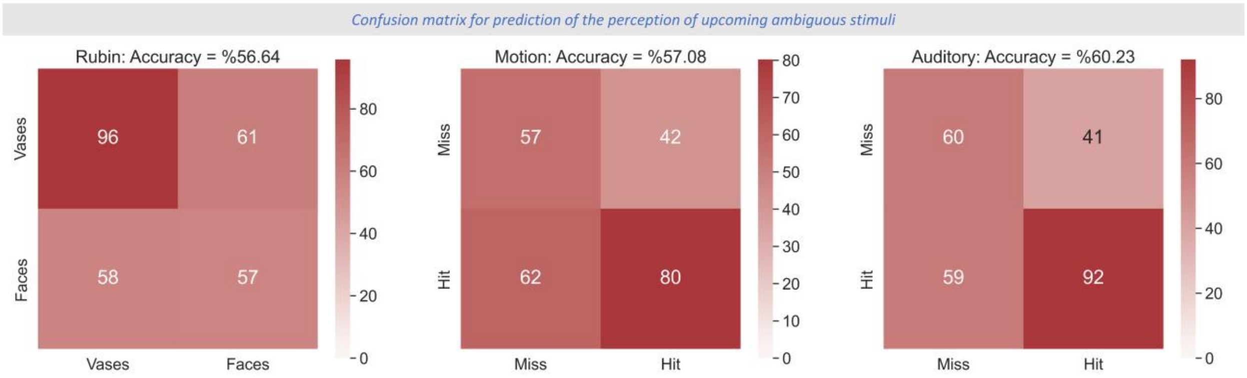


*Fig. S1 -* ***Confusion matrices across tasks.*** *Results are shown for Rubin, Motion detection, and Auditory vigilance tasks from left to right. In each confusion matrix, rows indicate true labels while columns indicate labels predicted by the trained model. The numbers in each cell show the number of trials with the corresponding predictions and true labels.*

### Cross-percept differences of the connectomes across all tasks

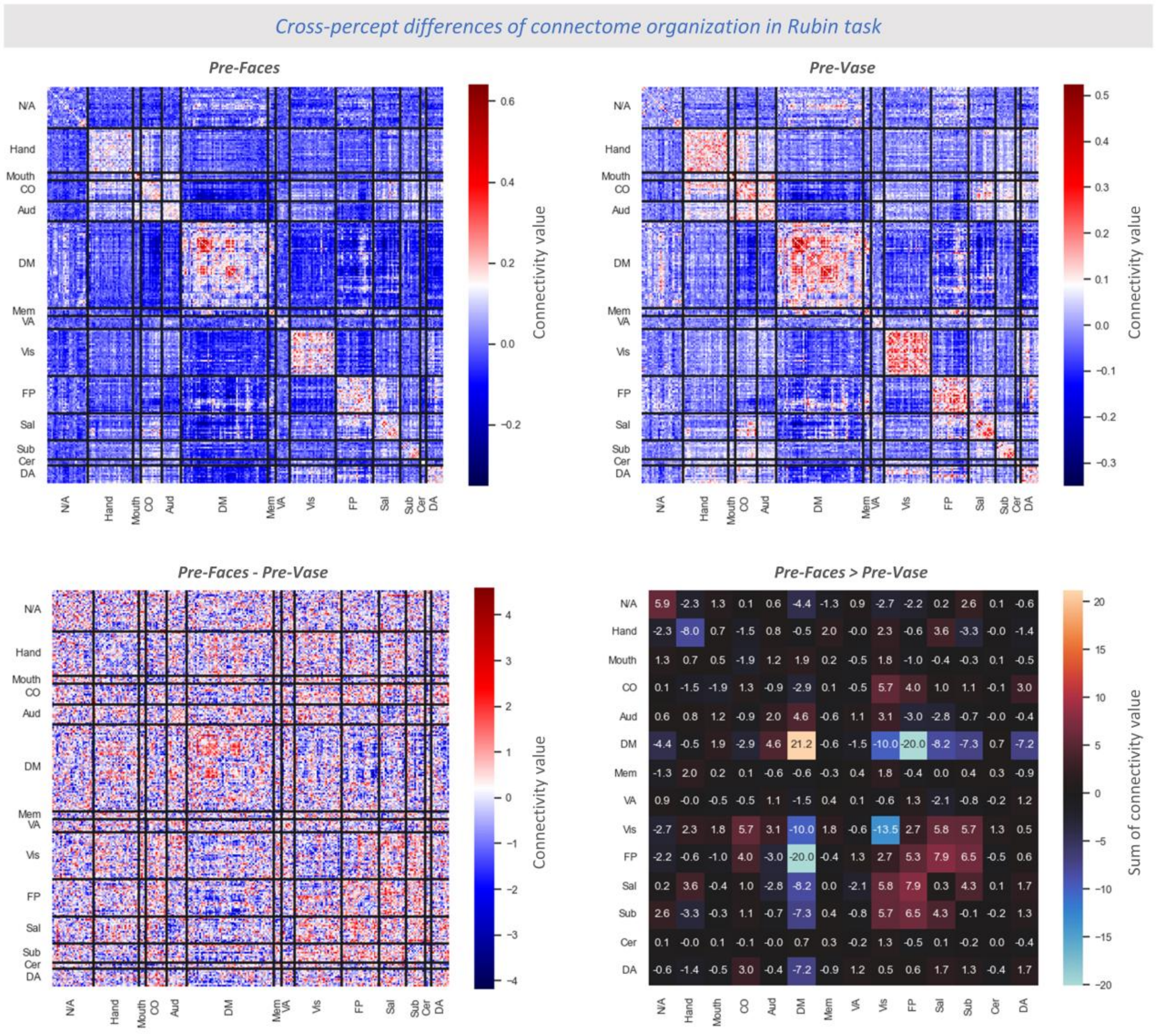


*Fig. S2 –* ***Cross-percept differences in connectome organization for Rubin task.*** *Top panels show the pre-stimulus connectome pattern averaged across all trials and timepoints. The top left panel corresponds to analytic target percept while the top right panel relates to analytic non-target percept. The bottom left panel depicts the differences between the top panels (target – non-target). The bottom right panel illustrates the sum of differences across all connections within every pair of canonical ICN.*

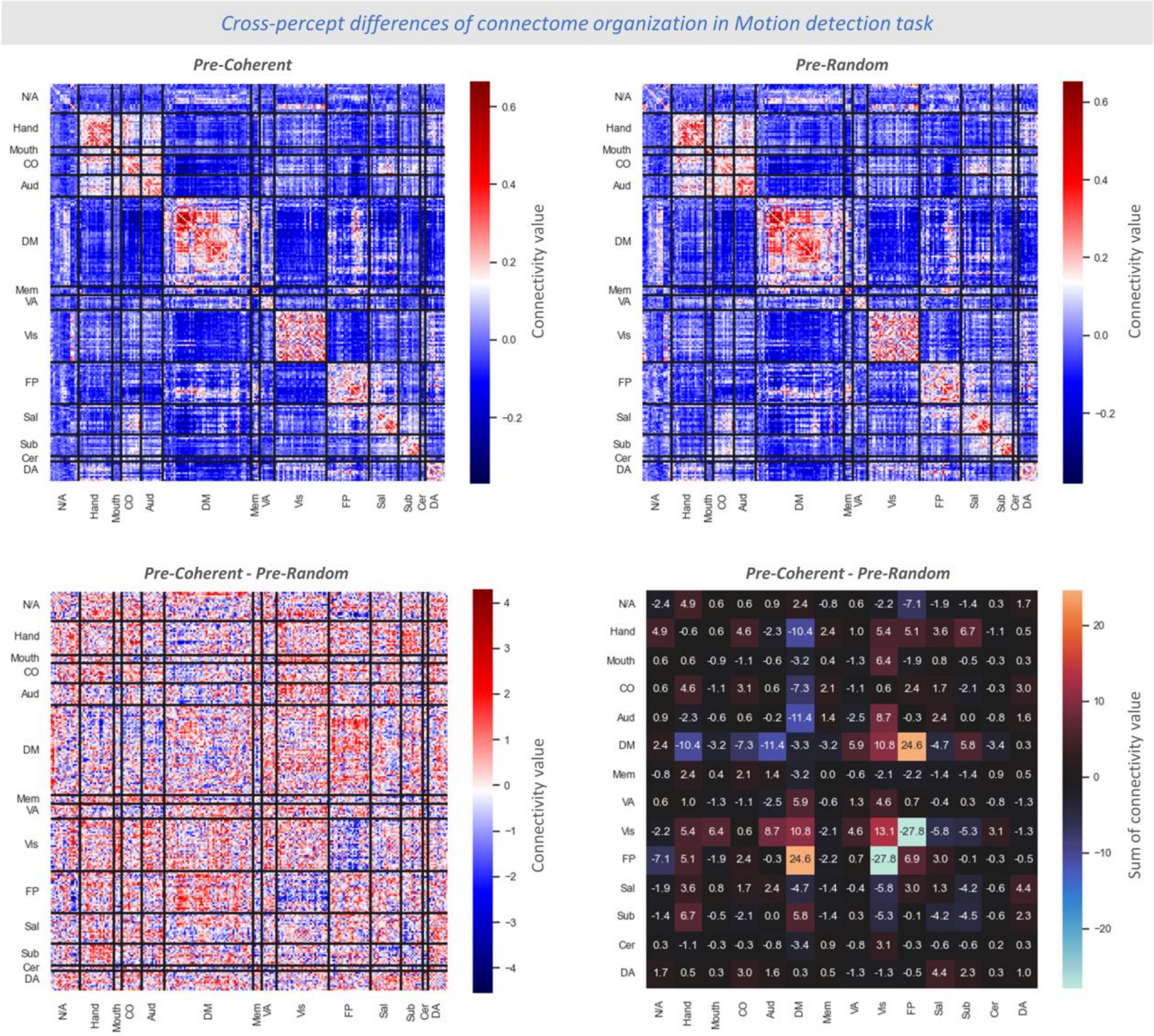


*Fig. S3* – ***Cross-percept differences in connectome organization for Motion detection task.***

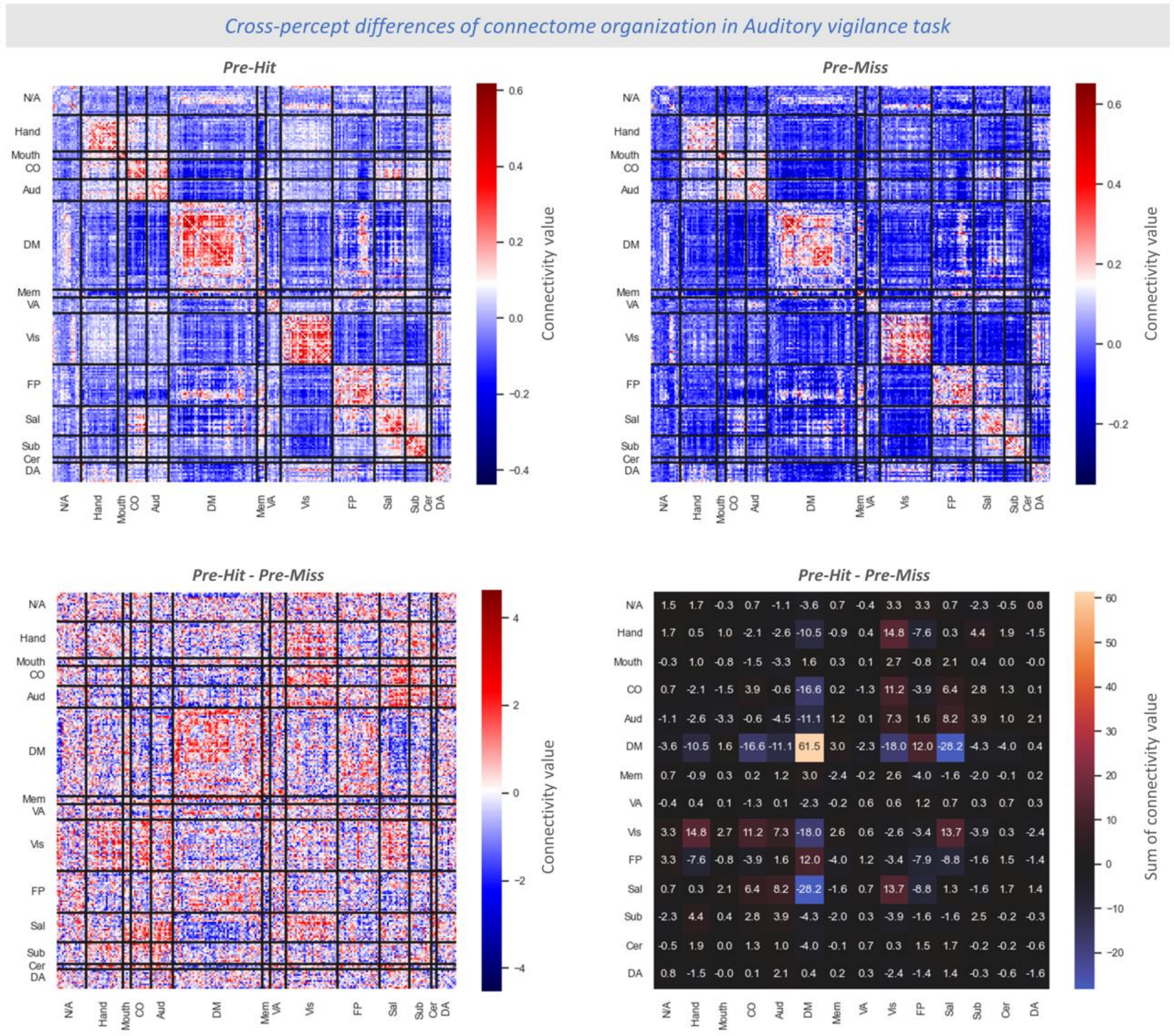


*Fig. S4 -* ***Cross-percept differences in connectome organization for Auditory vigilance task.***

### Investigation of demographic factors

We ensured that the task-specificity of predictive edges are not driven by the differences in cohorts' demographics across tasks. To this purpose, we added age and sex of the subjects of each task to our ANOVA analysis as covariate variables. Note that we excluded Rubin task from this complementary analysis since we did not have access to age and sex information of the individuals of this task. As shown in Fig. 4.15, the obtained $F$-values before and after adding the covariates for the two-level ANOVA (Motion detection and Auditory vigilance) showed only negligible changes across all connections ($r = 0.99$).

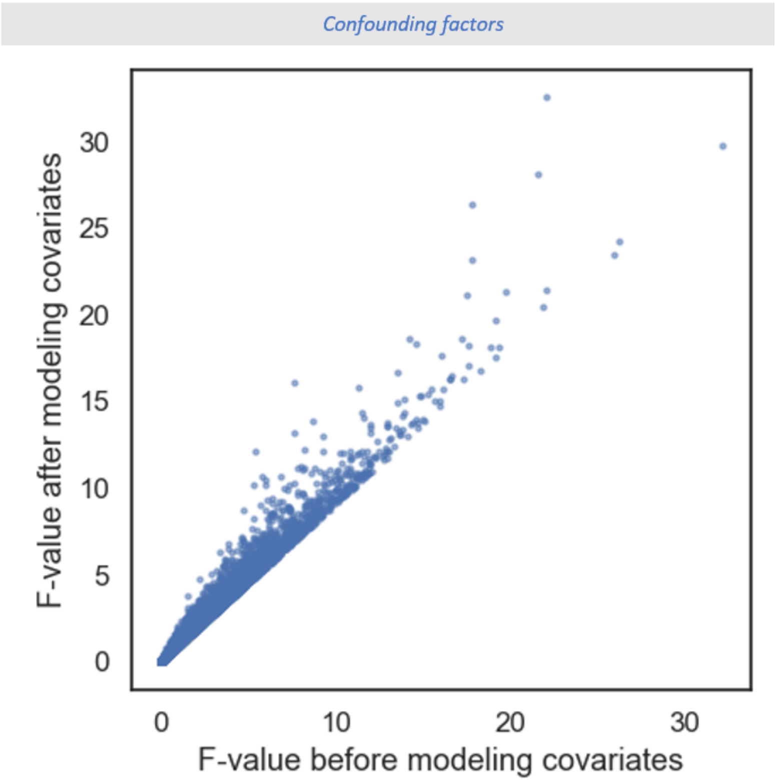


*Fig. S5 – **ANOVA F-values before and after modeling age and sex as covariates.** Horizontal and vertical axes respectively show the F-values of the ANOVA analysis before and after modeling the age and sex of participants as the model covariates, only for a Motion detection and Auditory vigilance comparison. The scores were highly correlated (r = 0.99) indicating that our findings of task-specific predictive connections are not driven by cross-task variations in sex and age of participant cohorts.*

**Perceptual effects of pre-stimulus connectivity versus regional activity**

The following analysis is analogous to the SVM approach in the main manuscript, with the exception that the available features comprised not only connection-wise connectivity values, but also region-wise activity levels. Thus, the total number of available features undergoing feature selection comprise *all* FC values plus 236 activity values, on which feature selection was performed. We did not observe an increase in model performance in any task (t-test across the 20 folds: $t_{19}$=-0.76, $p$=0.46 for Rubin; $t_{19}$=-1.36, $p$=0.19 for Motion; $t_{19}$=-1.12, $p$=0.28 for Auditory). Specifically, the prediction accuracy for Rubin, Motion, and Auditory tasks were respectively 0.57, 0.57, and 0.60 when the model was trained on FC values only, and 0.56, 0.56, and 0.59 when the model was trained on both FC and activity values.

Next, we compared the proportion of all available features among connectivity and activity values, respectively, that were selected by the feature selection procedure. In all tasks, a larger proportion of connectivity features than activity features were selected as predictive (t-tests for Rubin, Motion, and Auditory tasks: $t_{11}$ = 2.82; $p$ = 0.011, $t_{11}$ = 3.47; $p$ = 0.003, and $t_{10}$ = 7.59; $p$ = 0.000). This observation suggests that functional connectivity, impact perception of upcoming stimuli over and above regional activity alone.

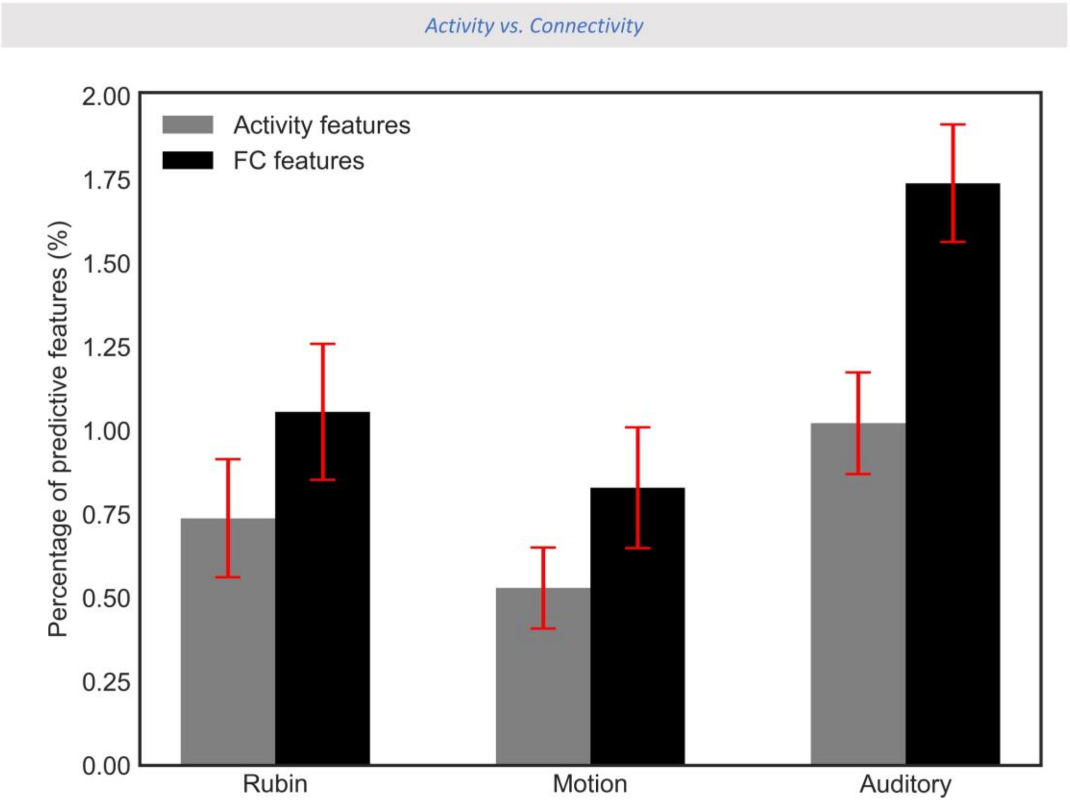


*Fig. S6 – **Percentage of within-region activity versus cross-region connectivity chosen during feature selection in each task.** Each columns set corresponds to a task. Within each set, gray or black bars indicate the percentage of predictive features respectively among regional activations and functional connectivity values. The percentage was calculated relative to the total number of features in the corresponding set. The error bar represents the standard error across twenty cross-validation folds of training data.*

### Global Network Topology

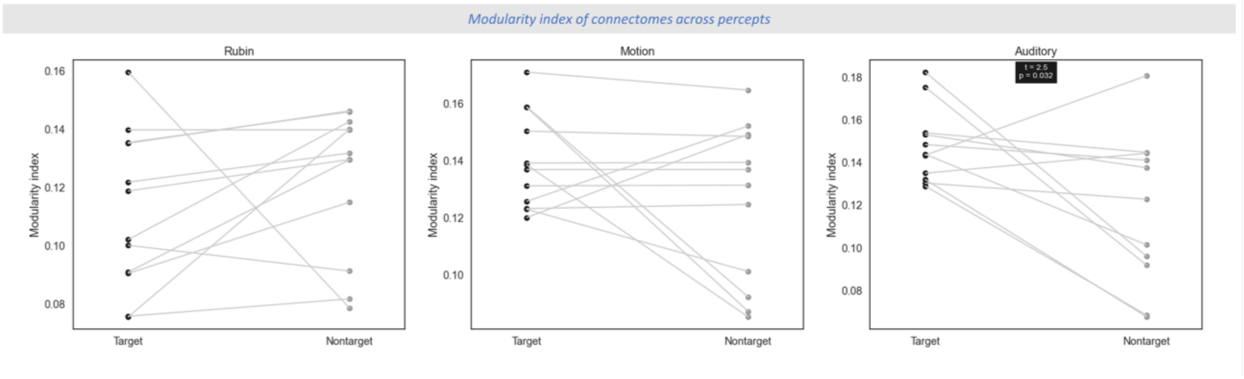


*Fig. S7 –* ***Comparison of Modularity index of pre-stimulus connectome across percepts in each task.*** *Overall Modularity index for every subject (shown as a filled circle) is shown across percepts (analytic target as black and analytic non-target as gray) in each task. From left to right, each subplot corresponds to Rubin, Motion detection, and Auditory vigilance task. Conditions with significant cross-percept differences are identified with an inset containing the statistical test results.*

*Table S1 – **Power-Petersen ROIs involving the predictive connections of Rubin task, including MNI coordinates and their corresponding resting state network.** Note that regions are sorted from top to bottom according to a counterclockwise order on the circular connectome view shown in Fig. 4 (Starting from Salience network, ending with Fronto-Parietal network).*

| ROI | X | Y | Z | Network | Avg_F | Edges |
|---|---|---|---|---|---|---|
| 25 | 28.54 | -39.24 | 59.17 | Hand | 5.9 | 1 |
| 40 | 3.45 | -17.44 | 58.45 | Hand | 6.2 | 1 |
| 44 | 51.14 | -5.8 | 32.42 | Mouth | 6.4 | 1 |
| 49 | 19.33 | -7.71 | 63.88 | CO | 7.4 | 1 |
| 58 | -51.26 | 8.26 | -2.06 | CO | 8.8 | 1 |
| 99 | -16.4 | 28.52 | 53.05 | DM | 7.6 | 1 |
| 108 | 8.8 | 54.23 | 3.45 | DM | 6.4 | 1 |
| 110 | 7.51 | 42.49 | -5.35 | DM | 7.8 | 2 |
| 119 | 64.8 | -30.55 | -8.7 | DM | 6.2 | 1 |
| 137 | -46.17 | 31.26 | -13.03 | DM | 5.9 | 1 |
| 138 | -9.88 | 10.95 | 66.61 | VA | 6.2 | 1 |
| 148 | 19.81 | -65.56 | 1.72 | Vis | 8.9 | 1 |
| 195 | -42.09 | -54.98 | 44.74 | FP | 8.6 | 1 |
| 203 | 10.51 | -38.54 | 50.02 | Sal | 6.2 | 1 |
| 210 | 36.89 | 32.35 | -2.24 | Sal | 8.0 | 1 |
| 220 | -39.12 | 50.79 | 17.38 | Sal | 8.7 | 1 |
| 222 | 6.3 | -23.68 | -0.42 | Sub | 8.8 | 1 |
| 223 | -1.77 | -13.05 | 11.82 | Sub | 7.4 | 1 |
| 230 | 23.26 | 10.19 | 1.46 | Sub | 8.7 | 1 |
| 231 | 28.52 | 0.82 | 4.01 | Sub | 7.6 | 1 |
| 236 | -56.47 | -50.48 | 9.92 | VA | 7.5 | 1 |
| 237 | -55.3 | -39.89 | 13.51 | VA | 8.9 | 1 |
| 257 | 46.09 | -58.93 | 3.93 | DA | 8.6 | 1 |

*Table S2 – **Power-Petersen ROIs involving the predictive connections of Motion task, including MNI coordinates and their corresponding resting state network.** Note that regions are sorted from top to bottom according to a counterclockwise order on the circular connectome view shown in Fig. 4 (Starting from Salience network, ending with Fronto-Parietal network).*

| ROI | X | Y | Z | Network | Avg_F | Edges |
|---|---|---|---|---|---|---|
| 17 | -6.9 | -20.59 | 65.21 | Hand | 7.5 | 1 |
| 20 | -53.52 | -22.54 | 43.1 | Hand | 8 | 1 |
| 33 | -45.1 | -31.85 | 46.63 | Hand | 9.9 | 1 |
| 39 | 2.4 | -27.94 | 60.15 | Hand | 6.4 | 1 |
| 49 | 19.33 | -7.71 | 63.88 | CO | 12.9 | 1 |
| 51 | -10.48 | -2.1 | 42.02 | CO | 9.8 | 1 |
| 60 | 35.83 | 10.32 | 1.18 | CO | 8.3 | 1 |
| 63 | 57.88 | -15.62 | 7.49 | Aud | 9.8 | 1 |
| 123 | 52.16 | -2.43 | -16.4 | DM | 8 | 1 |
| 135 | 11.27 | -66.01 | 42.09 | Mem | 8 | 1 |
| 143 | 17.53 | -46.86 | -9.88 | Vis | 8.8 | 1 |
| 150 | 26.93 | -59.37 | -9.36 | Vis | 6.4 | 1 |
| 151 | -15.02 | -72.42 | -7.68 | Vis | 8 | 1 |
| 152 | -17.87 | -68.03 | 4.81 | Vis | 8.2 | 1 |
| 168 | -40.21 | -88.44 | -6.19 | Vis | 8.2 | 1 |
| 173 | 36.51 | -81.16 | 1.2 | Vis | 5.8 | 1 |
| 174 | -43.93 | 1.8 | 45.7 | FP | 5.6 | 1 |
| 175 | 47.98 | 24.56 | 26.5 | FP | 12.9 | 1 |
| 179 | 58.31 | -52.79 | -13.61 | FP | 8.8 | 1 |
| 181 | 33.6 | 54.22 | -12.95 | FP | 7.5 | 1 |
| 192 | 43.93 | -52.95 | 46.95 | FP | 5.8 | 1 |
| 221 | 1.75 | -24.25 | 30.36 | Mem | 11.3 | 1 |
| 225 | 11.75 | -17.18 | 7.54 | Sub | 11.3 | 1 |
| 230 | 23.26 | 10.19 | 1.46 | Sub | 9.9 | 1 |
| 231 | 28.52 | 0.82 | 4.01 | Sub | 8.3 | 1 |
| 235 | 53.9 | -42.76 | 21.83 | VA | 5.3 | 1 |
| 243 | -16.31 | -65.28 | -19.69 | Cer | 5.3 | 1 |
| 264 | 28.56 | -4.62 | 53.99 | DA | 5.6 | 1 |

*Table S3 –* ***Power-Petersen ROIs involving the predictive connections of Auditory task, including MNI coordinates and their corresponding resting state network.*** *Note that regions are sorted from top to bottom according to a counterclockwise order on the circular connectome view shown in Fig. 4 (Starting from Salience network, ending with Fronto-Parietal network).*

| ROI | X | Y | Z | Network | Avg_F | Edges |
|---|---|---|---|---|---|---|
| 13 | -7.12 | -52.22 | 60.71 | Hand | 6.4 | 1 |
| 33 | -45.10 | -31.85 | 46.63 | Hand | 10.8 | 1 |
| 34 | -20.66 | -31.33 | 60.85 | Hand | 8.1 | 1 |
| 41 | 37.74 | -17.30 | 45.01 | Hand | 9.4 | 2 |
| 47 | -2.88 | 2.38 | 53.21 | CO | 15.1 | 1 |
| 49 | 19.33 | -7.71 | 63.88 | CO | 7.5 | 2 |
| 60 | 35.83 | 10.32 | 1.18 | CO | 6 | 1 |
| 63 | 57.88 | -15.62 | 7.49 | Aud | 8.1 | 1 |
| 68 | -49.77 | -34.36 | 25.74 | Aud | 8.2 | 1 |
| 72 | 59.40 | -17.34 | 28.69 | Aud | 15.1 | 1 |
| 86 | -44.45 | -64.64 | 34.78 | DM | 6.4 | 1 |
| 90 | -11.29 | -56.20 | 15.60 | DM | 6.3 | 1 |
| 102 | 12.73 | 54.87 | 38.19 | DM | 8.1 | 1 |
| 106 | 6.11 | 63.98 | 21.96 | DM | 8.1 | 1 |
| 123 | 52.16 | -2.43 | -16.40 | DM | 7.7 | 1 |
| 125 | 26.94 | -37.34 | -12.76 | DM | 9.3 | 1 |
| 135 | 11.27 | -66.01 | 42.09 | Mem | 5.8 | 1 |
| 144 | 39.98 | -72.49 | 14.36 | Vis | 8.7 | 1 |
| 145 | 8.45 | -71.84 | 10.79 | Vis | 10 | 1 |
| 148 | 19.81 | -65.56 | 1.72 | Vis | 8.8 | 2 |
| 157 | 28.68 | -76.62 | 25.42 | Vis | 9.3 | 1 |
| 162 | 24.41 | -87.21 | 24.01 | Vis | 8.3 | 1 |
| 173 | 36.51 | -81.16 | 1.20 | Vis | 10.8 | 1 |
| 177 | -52.60 | -48.83 | 42.50 | FP | 14.2 | 1 |
| 187 | -41.06 | 5.81 | 32.72 | FP | 8.7 | 1 |
| 188 | -42.23 | 38.21 | 21.35 | FP | 10 | 1 |
| 192 | 43.93 | -52.95 | 46.95 | FP | 7.7 | 1 |
| 201 | -42.10 | 24.68 | 29.53 | FP | 8.1 | 1 |
| 206 | 31.24 | 32.79 | 26.39 | Sal | 5.8 | 1 |
| 210 | 36.89 | 32.35 | -2.24 | Sal | 6.5 | 1 |
| 217 | 10.26 | 22.06 | 27.48 | Sal | 11.2 | 2 |
| 218 | 31.07 | 55.71 | 14.49 | Sal | 5.5 | 1 |
| 222 | 6.30 | -23.68 | -0.42 | Sub | 10.8 | 1 |
| 225 | 11.75 | -17.18 | 7.54 | Sub | 8.3 | 1 |
| 238 | 51.52 | -32.52 | 7.55 | VA | 6.5 | 1 |
| 241 | 52.68 | 32.58 | 0.57 | VA | 9.5 | 1 |
| 242 | -49.07 | 25.13 | -0.98 | VA | 10.8 | 1 |
| 245 | 22.43 | -57.55 | -23.11 | Cer | 9 | 1 |
| 258 | 25.34 | -58.18 | 60.34 | DA | 5.5 | 1 |
| 261 | -32.23 | -1.08 | 54.06 | DA | 12.5 | 1 |

*Table S4 –* ***A list of the 264 ROIs from the Power–Petersen atlas with their corresponding resting state network.*** *Regions labeled "N/A" were excluded from analyses but retained to preserve index order. The values in ROI column correspond across Tables S1-S4.*

| ROI | X | Y | Z | Network |
|---|---|---|---|---|
| 1 | -24.66 | -97.84 | -12.33 | N/A |
| 2 | 26.68 | -97.30 | -13.49 | N/A |
| 3 | 23.96 | 31.94 | -17.78 | N/A |
| 4 | -56.16 | -44.76 | -24.23 | N/A |
| 5 | 8.13 | 41.12 | -24.31 | N/A |
| 6 | -21.38 | -22.22 | -19.97 | N/A |
| 7 | 17.44 | -28.06 | -17.32 | N/A |
| 8 | -37.26 | -28.80 | -25.58 | N/A |
| 9 | 64.60 | -24.41 | -18.57 | N/A |
| 10 | 51.79 | -34.17 | -27.23 | N/A |
| 11 | 55.18 | -30.80 | -16.93 | N/A |
| 12 | 33.55 | 38.46 | -12.03 | N/A |
| 13 | -7.12 | -52.22 | 60.71 | Hand |
| 14 | -13.74 | -17.95 | 39.84 | Hand |
| 15 | 0.05 | -14.53 | 46.74 | Hand |
| 16 | 9.50 | -1.84 | 44.73 | Hand |
| 17 | -6.90 | -20.59 | 65.21 | Hand |
| 18 | -6.79 | -33.09 | 72.27 | Hand |
| 19 | 13.19 | -32.82 | 74.98 | Hand |
| 20 | -53.52 | -22.54 | 43.10 | Hand |
| 21 | 28.88 | -16.95 | 70.55 | Hand |
| 22 | 9.94 | -45.52 | 72.63 | Hand |
| 23 | -22.50 | -30.10 | 72.44 | Hand |
| 24 | -39.63 | -19.04 | 54.21 | Hand |
| 25 | 28.54 | -39.24 | 59.17 | Hand |
| 26 | 50.24 | -20.37 | 41.74 | Hand |
| 27 | -38.28 | -27.17 | 69.45 | Hand |
| 28 | 20.21 | -28.80 | 59.80 | Hand |
| 29 | 44.34 | -7.55 | 56.98 | Hand |
| 30 | -29.10 | -43.00 | 60.66 | Hand |
| 31 | 10.09 | -17.10 | 74.14 | Hand |
| 32 | 22.45 | -42.29 | 68.99 | Hand |
| 33 | -45.10 | -31.85 | 46.63 | Hand |
| 34 | -20.66 | -31.33 | 60.85 | Hand |
| 35 | -12.96 | -17.34 | 74.66 | Hand |
| 36 | 42.14 | -20.24 | 54.59 | Hand |
| 37 | -38.24 | -14.57 | 68.72 | Hand |
| 38 | -16.25 | -45.80 | 73.22 | Hand |
| 39 | 2.40 | -27.94 | 60.15 | Hand |
| 40 | 3.45 | -17.44 | 58.45 | Hand |
| 41 | 37.74 | -17.30 | 45.01 | Hand |
| 42 | -49.47 | -11.06 | 34.95 | Mouth |
| 43 | 36.04 | -9.44 | 13.95 | Mouth |
| 44 | 51.14 | -5.80 | 32.42 | Mouth |
| 45 | -52.84 | -10.23 | 24.41 | Mouth |
| 46 | 65.64 | -7.88 | 24.83 | Mouth |
| 47 | -2.88 | 2.38 | 53.21 | CO |
| 48 | 54.22 | -27.83 | 33.64 | CO |
| 49 | 19.33 | -7.71 | 63.88 | CO |
| 50 | -16.14 | -4.82 | 70.83 | CO |
| 51 | -10.48 | -2.10 | 42.02 | CO |
| 52 | 36.73 | 0.78 | -3.57 | CO |
| 53 | 13.21 | -1.36 | 69.98 | CO |
| 54 | 6.52 | 7.69 | 50.58 | CO |
| 55 | -44.76 | 0.10 | 8.83 | CO |
| 56 | 49.40 | 8.32 | -1.12 | CO |
| 57 | -34.37 | 3.29 | 4.19 | CO |
| 58 | -51.26 | 8.26 | -2.06 | CO |
| 59 | -5.33 | 17.80 | 34.41 | CO |
| 60 | 35.83 | 10.32 | 1.18 | CO |
| 61 | 31.75 | -26.33 | 12.91 | Aud |
| 62 | 65.43 | -33.20 | 19.97 | Aud |
| 63 | 57.88 | -15.62 | 7.49 | Aud |
| 64 | -38.43 | -33.34 | 16.98 | Aud |
| 65 | -60.48 | -25.22 | 13.82 | Aud |
| 66 | -49.14 | -26.30 | 5.18 | Aud |
| 67 | 43.45 | -22.93 | 19.85 | Aud |
| 68 | -49.77 | -34.36 | 25.74 | Aud |
| 69 | -52.92 | -21.83 | 22.97 | Aud |
| 70 | -55.22 | -9.42 | 11.73 | Aud |
| 71 | 55.96 | -5.03 | 13.25 | Aud |
| 72 | 59.40 | -17.34 | 28.69 | Aud |
| 73 | -30.12 | -27.02 | 12.20 | Aud |
| 74 | -40.50 | -75.27 | 25.80 | DM |
| 75 | 5.55 | 66.69 | -3.55 | DM |
| 76 | 8.36 | 47.59 | -15.18 | DM |
| 77 | -12.60 | -39.64 | 0.93 | DM |
| 78 | -17.65 | 63.19 | -9.17 | DM |
| 79 | -45.79 | -60.69 | 20.85 | DM |
| 80 | 43.43 | -72.21 | 28.00 | DM |
| 81 | -43.58 | 11.99 | -34.15 | DM |
| 82 | 45.64 | 16.20 | -30.02 | DM |
| 83 | -68.47 | -22.66 | -15.74 | DM |
| 84 | -57.97 | -25.69 | -14.73 | N/A |
| 85 | 27.06 | 16.22 | -16.93 | N/A |
| 86 | -44.45 | -64.64 | 34.78 | DM |
| 87 | -39.05 | -74.95 | 43.72 | DM |
| 88 | -6.84 | -54.90 | 27.05 | DM |
| 89 | 5.91 | -58.82 | 35.45 | DM |
| 90 | -11.29 | -56.20 | 15.60 | DM |
| 91 | -2.94 | -48.79 | 12.87 | DM |
| 92 | 7.94 | -48.37 | 30.57 | DM |
| 93 | 15.12 | -63.09 | 25.98 | DM |
| 94 | -2.20 | -36.68 | 43.85 | DM |
| 95 | 10.77 | -53.83 | 17.09 | DM |
| 96 | 52.04 | -59.37 | 35.52 | DM |
| 97 | 23.33 | 33.07 | 47.68 | DM |
| 98 | -10.09 | 39.09 | 52.29 | DM |
| 99 | -16.40 | 28.52 | 53.05 | DM |
| 100 | -35.36 | 19.86 | 50.80 | DM |
| 101 | 22.11 | 39.21 | 38.90 | DM |
| 102 | 12.73 | 54.87 | 38.19 | DM |
| 103 | -10.33 | 54.63 | 38.71 | DM |
| 104 | -19.78 | 45.07 | 39.48 | DM |
| 105 | 5.94 | 54.42 | 16.18 | DM |
| 106 | 6.11 | 63.98 | 21.96 | DM |
| 107 | -7.04 | 50.82 | -1.29 | DM |
| 108 | 8.80 | 54.23 | 3.45 | DM |
| 109 | -3.06 | 44.41 | -9.46 | DM |
| 110 | 7.51 | 42.49 | -5.35 | DM |
| 111 | -11.06 | 44.62 | 7.61 | DM |
| 112 | -2.06 | 37.85 | 36.34 | DM |
| 113 | -2.50 | 41.70 | 16.05 | DM |
| 114 | -20.16 | 63.65 | 19.39 | DM |
| 115 | -7.55 | 48.08 | 23.18 | DM |
| 116 | 64.64 | -11.80 | -19.30 | DM |
| 117 | -55.72 | -12.96 | -10.24 | DM |
| 118 | -57.75 | -29.70 | -3.94 | DM |
| 119 | 64.80 | -30.55 | -8.70 | DM |
| 120 | -68.30 | -41.41 | -5.14 | DM |
| 121 | 13.08 | 29.99 | 58.65 | DM |
| 122 | 12.25 | 35.63 | 20.30 | DM |
| 123 | 52.16 | -2.43 | -16.40 | DM |
| 124 | -26.44 | -39.95 | -8.26 | DM |
| 125 | 26.94 | -37.34 | -12.76 | DM |
| 126 | -33.93 | -38.06 | -15.60 | DM |
| 127 | 28.46 | -76.56 | -31.64 | DM |
| 128 | 51.90 | 6.81 | -29.61 | DM |
| 129 | -52.89 | 2.55 | -27.06 | DM |
| 130 | 46.68 | -50.08 | 28.76 | DM |
| 131 | -49.30 | -42.15 | 0.83 | DM |
| 132 | -30.63 | 18.71 | -18.98 | N/A |
| 133 | -2.47 | -34.80 | 31.07 | Mem |
| 134 | -6.58 | -71.47 | 41.74 | Mem |
| 135 | 11.27 | -66.01 | 42.09 | Mem |
| 136 | 4.20 | -48.06 | 50.71 | Mem |
| 137 | -46.17 | 31.26 | -13.03 | DM |
| 138 | -9.88 | 10.95 | 66.61 | VA |
| 139 | 49.26 | 35.47 | -12.20 | DM |
| 140 | 7.98 | -91.08 | -7.10 | N/A |
| 141 | 17.27 | -91.09 | -13.64 | N/A |
| 142 | -12.08 | -94.56 | -12.80 | N/A |
| 143 | 17.53 | -46.86 | -9.88 | Vis |
| 144 | 39.98 | -72.49 | 14.36 | Vis |
| 145 | 8.45 | -71.84 | 10.79 | Vis |
| 146 | -8.43 | -80.50 | 7.44 | Vis |
| 147 | -28.07 | -79.45 | 19.43 | Vis |
| 148 | 19.81 | -65.56 | 1.72 | Vis |
| 149 | -23.94 | -90.98 | 18.96 | Vis |
| 150 | 26.93 | -59.37 | -9.36 | Vis |
| 151 | -15.02 | -72.42 | -7.68 | Vis |
| 152 | -17.87 | -68.03 | 4.81 | Vis |
| 153 | 42.52 | -78.17 | -11.78 | Vis |
| 154 | -46.54 | -75.95 | -9.95 | Vis |
| 155 | -14.22 | -90.66 | 31.40 | Vis |
| 156 | 15.27 | -87.09 | 36.89 | Vis |
| 157 | 28.68 | -76.62 | 25.42 | Vis |
| 158 | 19.64 | -85.62 | -2.39 | Vis |
| 159 | 15.18 | -76.68 | 31.00 | Vis |
| 160 | -15.85 | -52.34 | -1.43 | Vis |
| 161 | 41.60 | -65.50 | -8.27 | Vis |
| 162 | 24.41 | -87.21 | 24.01 | Vis |
| 163 | 5.59 | -71.65 | 23.52 | Vis |
| 164 | -42.10 | -73.62 | 0.38 | Vis |
| 165 | 25.66 | -79.47 | -15.56 | Vis |
| 166 | -16.21 | -76.97 | 33.82 | Vis |
| 167 | -2.88 | -81.25 | 21.10 | Vis |
| 168 | -40.21 | -88.44 | -6.19 | Vis |
| 169 | 36.76 | -84.11 | 12.99 | Vis |
| 170 | 6.21 | -81.41 | 6.11 | Vis |
| 171 | -26.39 | -90.23 | 3.12 | Vis |
| 172 | -33.00 | -79.02 | -13.24 | Vis |
| 173 | 36.51 | -81.16 | 1.20 | Vis |
| 174 | -43.93 | 1.80 | 45.70 | FP |
| 175 | 47.98 | 24.56 | 26.50 | FP |
| 176 | -46.50 | 10.85 | 23.04 | FP |
| 177 | -52.60 | -48.83 | 42.50 | FP |
| 178 | -22.53 | 10.76 | 63.73 | FP |
| 179 | 58.31 | -52.79 | -13.61 | FP |
| 180 | 24.07 | 44.61 | -15.35 | FP |
| 181 | 33.60 | 54.22 | -12.95 | FP |
| 182 | -21.14 | 40.87 | -20.48 | N/A |
| 183 | -17.51 | -75.89 | -24.33 | N/A |
| 184 | 16.85 | -79.89 | -34.36 | N/A |
| 185 | 34.72 | -67.08 | -34.45 | N/A |
| 186 | 47.01 | 9.93 | 32.66 | FP |
| 187 | -41.06 | 5.81 | 32.72 | FP |
| 188 | -42.23 | 38.21 | 21.35 | FP |
| 189 | 38.37 | 43.18 | 15.06 | FP |
| 190 | 49.18 | -42.41 | 45.16 | FP |
| 191 | -28.40 | -57.93 | 47.78 | FP |
| 192 | 43.93 | -52.95 | 46.95 | FP |
| 193 | 31.83 | 14.37 | 55.98 | FP |
| 194 | 37.45 | -64.70 | 40.38 | FP |
| 195 | -42.09 | -54.98 | 44.74 | FP |
| 196 | 39.87 | 18.39 | 39.72 | FP |
| 197 | -34.16 | 54.83 | 4.36 | FP |
| 198 | -41.68 | 45.16 | -2.31 | FP |
| 199 | 33.38 | -53.12 | 44.02 | FP |
| 200 | 43.25 | 49.25 | -2.31 | FP |
| 201 | -42.10 | 24.68 | 29.53 | FP |
| 202 | -2.98 | 26.41 | 44.42 | FP |
| 203 | 10.51 | -38.54 | 50.02 | Sal |
| 204 | 55.27 | -44.59 | 36.70 | Sal |
| 205 | 42.05 | -0.39 | 47.10 | Sal |
| 206 | 31.24 | 32.79 | 26.39 | Sal |
| 207 | 47.60 | 22.16 | 9.74 | Sal |
| 208 | -35.44 | 20.03 | 0.07 | Sal |
| 209 | 35.91 | 21.91 | 2.62 | Sal |
| 210 | 36.89 | 32.35 | -2.24 | Sal |
| 211 | 33.56 | 16.45 | -7.58 | Sal |
| 212 | -10.76 | 25.99 | 24.54 | Sal |
| 213 | -0.94 | 14.86 | 43.99 | Sal |
| 214 | -27.50 | 52.04 | 21.28 | Sal |
| 215 | -0.20 | 30.35 | 27.22 | Sal |
| 216 | 5.23 | 23.22 | 37.03 | Sal |
| 217 | 10.26 | 22.06 | 27.48 | Sal |
| 218 | 31.07 | 55.71 | 14.49 | Sal |
| 219 | 26.07 | 49.56 | 26.58 | Sal |
| 220 | -39.12 | 50.79 | 17.38 | Sal |
| 221 | 1.75 | -24.25 | 30.36 | Mem |
| 222 | 6.30 | -23.68 | -0.42 | Sub |
| 223 | -1.77 | -13.05 | 11.82 | Sub |
| 224 | -10.28 | -18.48 | 7.04 | Sub |
| 225 | 11.75 | -17.18 | 7.54 | Sub |
| 226 | -5.33 | -28.08 | -4.13 | Sub |
| 227 | -21.97 | 7.48 | -4.78 | Sub |
| 228 | -15.41 | 3.57 | 7.99 | Sub |
| 229 | 30.50 | -13.92 | 1.65 | Sub |
| 230 | 23.26 | 10.19 | 1.46 | Sub |
| 231 | 28.52 | 0.82 | 4.01 | Sub |
| 232 | -31.38 | -11.48 | -0.30 | Sub |
| 233 | 14.98 | 4.94 | 7.24 | Sub |
| 234 | 8.62 | -3.57 | 5.76 | Sub |
| 235 | 53.90 | -42.76 | 21.83 | VA |
| 236 | -56.47 | -50.48 | 9.92 | VA |
| 237 | -55.30 | -39.89 | 13.51 | VA |
| 238 | 51.52 | -32.52 | 7.55 | VA |
| 239 | 51.28 | -28.52 | -4.30 | VA |
| 240 | 55.75 | -46.07 | 11.42 | VA |
| 241 | 52.68 | 32.58 | 0.57 | VA |
| 242 | -49.07 | 25.13 | -0.98 | VA |
| 243 | -16.31 | -65.28 | -19.69 | Cer |
| 244 | -32.12 | -55.03 | -25.22 | Cer |
| 245 | 22.43 | -57.55 | -23.11 | Cer |
| 246 | 0.51 | -61.91 | -18.14 | Cer |
| 247 | 32.85 | -12.41 | -34.41 | N/A |
| 248 | -31.13 | -9.99 | -36.32 | N/A |
| 249 | 48.52 | -2.85 | -38.49 | N/A |
| 250 | -50.06 | -7.09 | -39.24 | N/A |
| 251 | 9.61 | -61.50 | 60.88 | DA |
| 252 | -52.44 | -63.14 | 5.29 | DA |
| 253 | -46.68 | -50.91 | -20.91 | N/A |
| 254 | 45.68 | -46.67 | -16.85 | N/A |
| 255 | 47.21 | -29.75 | 48.70 | Hand |
| 256 | 21.90 | -64.74 | 48.12 | DA |
| 257 | 46.09 | -58.93 | 3.93 | DA |
| 258 | 25.34 | -58.18 | 60.34 | DA |
| 259 | -32.56 | -46.42 | 47.20 | DA |
| 260 | -26.60 | -70.72 | 36.86 | DA |
| 261 | -32.23 | -1.08 | 54.06 | DA |
| 262 | -42.26 | -60.12 | -8.85 | DA |
| 263 | -16.50 | -58.57 | 64.46 | DA |
| 264 | 28.56 | -4.62 | 53.99 | DA |
